# Neural correlates of location–response compatibility in an immersive virtual-reality Attention Network Test: a multiverse electroencephalography analysis

**DOI:** 10.64898/2026.08.20.746062

**Authors:** David Levi Tekampe, Philip S. Santangelo, Anxhela Sulaj, Philipp Tekampe, Michael Schwartze, Lars Hausfeld

## Abstract

Immersive virtual reality (VR) can preserve the logic of laboratory attention tasks while altering the perceptual-action context in which attentional control is expressed. In this study, we examined the neural underpinnings of location-response compatibility in a VR adaptation of the Attention Network Test-Revised (ANT-VR), using a restricted preprocessing multiverse to account for uncertainty arising from defensible EEG analysis choices. Forty-four young adults contributed complete ANT-VR behavioural data. Target-locked EEG analyses were conducted across 192 preprocessing branches, with branch-level participant contributions varying after quality check and trial-count filtering. The contrast compared location-response incompatible with compatible trials and was balanced within participants across cue-target interval, cue condition, flanker congruency, and, for spatial-cue trials, conditions in which spatial cues were valid or invalid for subsequent targets. Across the multiverse, the N2pc-window posterior-lateralisation contrast could be defined in all branches and showed high directional stability: the median incompatible-minus-compatible effect was 0.30 µV [IQR: 0.22 to 0.41], with positive effects in 192/192 branches, nominal evidence in 79/192 branches, and Holm-corrected evidence in 42/192 branches. Comparison across ERP measures indicated that this pattern was more consistent for N2pc-window posterior lateralisation than for P1, posterior N1, frontocentral N2, P3, or response-referenced C3/C4 measures. Comparison across C3/C4 reference frames further constrained the interpretation: target-location-referenced C3/C4 showed the strongest effect, whereas response-referenced C3/C4 was weaker. Behavioural analyses showed no reliable compatibility differences. These findings suggest that location-response compatibility in immersive ANT-VR modulates target-locked lateralised neural activity associated with spatial selection and target-location coding, rather than producing broad sensory, conflict-related, P3-related, or specifically response-referenced modulation.

**Significance Statement:** Behavioural performance can conceal neural effects of spatial compatibility, particularly in immersive tasks where perception and action are closely coupled. In the Attention Network Test–VR, location-response incompatibility produced no reliable reaction-time or accuracy cost, yet target-locked EEG showed a directionally stable posterior-lateralisation shift in the N2pc window across 192 defensible preprocessing specifications. Target-location-referenced C3/C4 activity showed stronger and more consistent modulation than response-referenced activity, arguing against a purely response-preparation account. These findings indicate that task-irrelevant target location can alter spatial-selection or target-location coding even when behavioural performance is unchanged, while illustrating how multiverse analysis can test whether such neural conclusions depend on preprocessing choices.

## 1. Introduction

Attention supports adaptive behaviour by prioritising relevant information and coordinating selection and control processes (Posner & Petersen, 1990; Petersen & Posner, 2012). The attention-network model distinguishes alerting, orienting, and executive control as partially dissociable but interacting attentional functions (Petersen & Posner, 2012). Behavioural performance reflects the combined outcome of multiple processing operations, potentially including perceptual and spatial selection, conflict monitoring, decision-related processing, and response selection and preparation (Eimer, 1998; Folstein & Van Petten, 2008; Hillyard & Anllo-Vento, 1998; Polich, 2007). Therefore, a small or absent behavioural cost does not imply that the underlying processing dynamics are free of competition; compatibility-related differences may be resolved, compensated for, or expressed neurally before becoming visible in reaction time or accuracy.

The original Attention Network Test combines cueing and flanker conflict (Fan et al., 2002), whereas the Attention Network Test-Revised (ANT-R) additionally incorporates cue-validity, cue-target-interval, and location-congruency manipulations (Fan et al., 2009). Spatial cues indicate the likely target location and can therefore be valid or invalid for the subsequent target. Separately, location-response compatibility refers to whether the physical target location corresponds to the required response side. In ANT-VR (Tekampe et al., 2023, 2026), target identity determines the required response side, whereas target location can be compatible or incompatible with that response. This object-based implementation of location-response compatibility is related to spatial stimulus-response compatibility and Simon-effect accounts, in which task-irrelevant spatial information influences response selection (Simon & Rudell, 1967; Kornblum et al., 1990; Lu & Proctor, 1995). However, the ANT-VR manipulation is not a canonical Simon effect because meaningful objects are embedded in an immersive scene and responses are made using handheld controllers.

Compared with non-immersive display formats, immersive VR introduces distinct three-dimensional, sensorimotor, presence-related, and embodiment-related cues (Bohil et al., 2011; Kilteni et al., 2012; Parsons, 2015). Immersive virtual reality changes more than the display medium. Relative to desktop tasks, VR can alter depth structure, spatial presence, self-location, embodiment, and controller-mediated action (Bohil et al., 2011; Kilteni et al., 2012; Parsons, 2015). Previous ANT-VR behavioural work indicates that immersive administration can preserve major ANT-R contrast patterns while moderating selected orienting- and conflict-related effects (Tekampe et al., 2023, 2026). However, previous ANT-VR analyses did not reveal a reliable behavioural location-response compatibility effect. The present EEG analysis therefore extends these behavioural findings by providing a temporally resolved account of the processing operations associated with location-response compatibility, allowing compatibility-related modulation to be characterised beyond the behavioural endpoint.

Event-related potentials provide temporally specific indices of candidate processing stages. P1 or posterior N1 modulation would be consistent with compatibility effects arising at relatively early visual processing stages, whereas N2pc modulation would indicate altered lateralised target prioritisation, spatial selection, or coding of the lateralised object (Luck & Hillyard, 1994; Eimer, 1996; Kiss et al., 2008). Frontocentral N2 or P3 modulation would point towards broader conflict-related, executive, or decisional consequences (Folstein & Van Petten, 2008; Polich, 2007). Lateralised C3/C4 activity further constrains interpretation with respect to reference frame: response-referenced effects may suggest response-specific activation, whereas target-location-referenced effects would argue against a simple lateralised-readiness-potential-like interpretation (Eimer, 1998; Praamstra & Oostenveld, 2003; Praamstra, 2007).

The present study examined the neural processes associated with location-response compatibility in ANT-VR and whether the resulting conclusions remained stable across defensible preprocessing specifications. We expected location-response compatibility to modulate target-locked lateralised EEG activity most clearly in the N2pc time window if the manipulation primarily affected spatial selection or target-location coding. In contrast, stronger P1 or posterior N1 effects would support early sensory modulation, stronger frontocentral N2 or P3 effects would support broader conflict-related or decisional consequences, and stronger response-referenced C3/C4 effects would support a more response-preparation-centred account. To assess whether the interpretation depended on particular but defensible EEG preprocessing choices, these contrasts were evaluated across a restricted preprocessing multiverse (Clayson et al., 2021).

## 2. Materials and methods

### 2.1. Participants and procedure

The EEG analyses used data from the ANT-VR cohort whose behavioural findings have been reported previously (Tekampe et al., 2026). Fifty young adult volunteers were recruited at Maastricht University, the Netherlands, between April 2022 and September 2023 through the Research Participation System, word of mouth, and local advertisements. Eligibility required normal or corrected-to-normal vision, normal hearing, and no self-reported history of a diagnosed neuropsychological disorder. All participants provided written informed consent. The study was approved by the Ethics Review Committee Psychology and Neuroscience at Maastricht University (249_29_02_2022; 23 March 2022). Procedures conformed to the Declaration of Helsinki.

Data from three participants were excluded because of technical issues during acquisition, leaving 47 participants with EEG data available for preprocessing. After protocol, behavioural-quality, and technical exclusions, the sample overlapping with the final behavioural cohort comprised 44 participants aged 18-31 years (M = 21.91, SD = 3.57), including 22 cisgender women, 21 cisgender men, and one non-binary participant. Thirty-nine participants were right-handed and five were left-handed. Final EEG sample sizes varied across preprocessing branches and contrasts because the analyses required branch × participant datasets that passed preprocessing and quality check (QC), overlapped with the behavioural sample, contained the required retained epochs, involved no interpolation of C3 or C4, and met the minimum trial count after cell-count filtering.

Participants completed practice trials before the experimental task and were instructed to respond quickly and accurately while minimising unnecessary movement. The full session included both a computerised ANT-R and the ANT-VR, with task order counterbalanced across participants. The present study reports the ANT-VR EEG analyses, focusing on the neural correlates of location-response compatibility.

### 2.2. ANT-VR task and compatibility contrast

The Virtual Reality Attention Network Task was administered using a Meta Quest 2 head-mounted display in standalone mode. Responses were made using the left and right VR controllers. The task retained the temporal structure of the Attention Network Test-Revised (ANT-R; Fan et al., 2009) while replacing arrow stimuli with everyday objects. For a visual depiction of the ANT-VR environment and stimulus configuration, see Tekampe et al., 2026. On each trial, participants viewed a virtual computer screen within an apartment-like virtual environment. The central target was either a phone or a tablet and was flanked by four congruent or incongruent objects. A phone and a tablet were also displayed laterally as response-mapping cues. Participants identified the central target object using the corresponding controller button.

Each trial consisted of a cue phase, a cue-target interval, a target/ flanker display, a response window, and an inter-trial interval. The cue condition was no cue, double cue, or spatial cue. Spatial cues were valid or invalid with respect to the subsequent target location. Cue-target intervals were 0, 400, or 800 ms. The target/flanker display lasted 500 ms, and responses were accepted for up to 1,700 ms after target onset. The interval from target offset to the next trial onset ranged from 2,000 to 12,000 ms.

The manipulation of interest was location-response compatibility. On compatible trials, the physical target location corresponded to the response side defined by target identity. On incompatible trials, the target location and target-defined response side were opposite. The EEG compatibility effect was defined as the incompatible-minus-compatible contrast. This contrast was estimated for each combination of flanker congruency, cue condition, cue-target interval, and spatial-cue validity for spatial-cue trials.

### 2.3. EEG acquisition and event processing

EEG was recorded using a mobile mBrainTrain SMARTING system (Belgrade, Serbia) and a 24-channel SMARTINGmobi cap with Ag/AgCl electrodes positioned according to the international 10-20 system. The mobile system allowed EEG to be recorded while preserving the immersive, controller-based ANT-VR setup used for behavioural testing. Signals were digitised at 250 Hz with 24-bit resolution. The common mode sense reference and driven right leg ground electrodes were positioned at FCz and Fpz, respectively. Three gyroscope channels were retained for movement auditing and movement-sensitive epoch rejection but were excluded from EEG filtering, referencing, independent component analysis (ICA), feature extraction, and statistical analyses.

ANT-VR event markers were streamed using Lab Streaming Layer (Kothe et al., 2025) and synchronised with the electroencephalographic recording using Smarting Streamer software (mBrainTrain, Belgrade, Serbia). The markers encoded trial phase, cue condition, cue-target interval, target location, target identity, target-defined response side, flanker congruency, and button responses. They were decoded using a study-specific MATLAB/ EEGLAB pipeline to generate branch-specific trial tables and epoch metadata. Before EEG feature extraction, we checked event timing, trial-phase order, target-to-response timing, duplicated or missing response markers, and agreement between the decoded trial tables and epoch metadata. Target and response markers served as the temporal references for target-locked and response-locked epoching, respectively. Because marker latency was not independently validated, interpretation focused on mean amplitudes within predefined component windows rather than precise single-sample latency estimates.

### 2.4. Restricted preprocessing multiverse

Several defensible preprocessing choices were available, including alternative approaches to line-noise attenuation, bad-channel handling, and referencing, which can influence subsequent EEG estimates (Bigdely-Shamlo et al., 2015). To assess whether conclusions concerning the location-response compatibility effect depended on a single preprocessing specification, we implemented a restricted preprocessing multiverse. EEG preprocessing was performed in MATLAB using EEGLAB-compatible routines.

The final multiverse comprised 192 preprocessing branches formed by crossing high-pass filter (0.1 vs. 0.5 Hz), low-pass filter (30 vs. 40 Hz), line-noise attenuation using CleanLine (Mullen, 2012) or a 50-Hz notch filter, artefact subspace reconstruction (ASR vs. no ASR; Chang et al., 2020), reference policy (average reference vs. linked mastoids), ICLabel-based component-rejection policy (conservative, moderate, or eye/muscle-only; Pion-Tonachini et al., 2019), and cleaning-stringency bundle. The cleaning-stringency bundle had two levels: canonical epoch rejection with a −200 to 0 ms target baseline and no movement-sensitive rejection, or strict epoch rejection with a −100 to 0 ms target baseline and gyro-sensitive rejection. The specification space therefore formed a balanced 2 × 2 × 2 × 2 × 2 × 3 × 2 design. The preprocessing factors and their branch-level outcome mapping are visualised in Figure 1.

**Figure 1.**
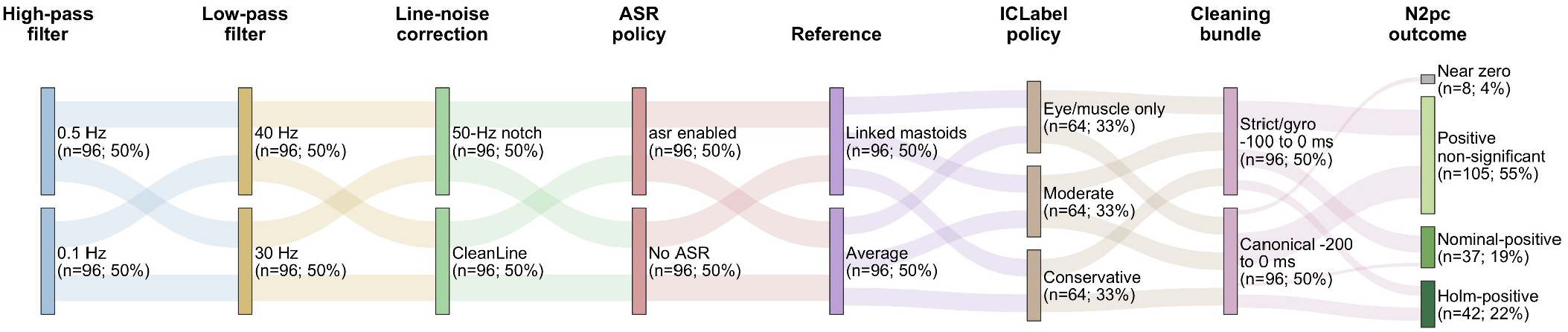
Restricted preprocessing multiverse and N2pc outcome mapping **Note.** Alluvial diagram showing the 192 preprocessing specifications in the restricted ANT-VR EEG multiverse and their classification according to the target-locked N2pc compatibility contrast, defined as incompatible minus compatible. Branches crossed high-pass filter, low-pass filter, line-noise attenuation, ASR policy, reference policy, ICLabel rejection policy, and cleaning-stringency bundle. Flow widths indicate the number of preprocessing specifications passing through each decision level and outcome category. Outcome categories show the N2pc result for each branch, with “near zero” denoting an absolute N2pc effect < 0.10 µV after significance categories were assigned. Because all branches were based on the same participants and raw data, they represent robustness across preprocessing choices rather than independent replications. ASR = artefact subspace reconstruction; N2pc = N2 posterior contralateral component.

The branches were treated as a bounded set of defensible preprocessing specifications for this ANT-VR EEG dataset. They do not represent all possible EEG pipelines, and the aim was not to identify an optimal branch. Rather, the multiverse quantified whether the compatibility effect and its interpretation were stable across plausible preprocessing alternatives. Because all branches reused the same participants, raw recordings, task labels, feature definitions, and statistical tests, branch-wise results were interpreted as robustness summaries rather than independent replications.

Raw EEG was imported from XDF files, channel locations were assigned, and FCz was appended as a channel-location/reference label rather than treated as a recorded electrode. Branch-specific line-noise correction, filtering, ASR policy, and referencing were then applied. Bad-channel handling followed a staged conservative policy: the prespecified T7 decision and automatically detected bad channels were logged, temporarily interpolated for ASR when necessary, excluded from pre-ICA referencing, ICA estimation, and epoch artefact rejection, and interpolated only after IC removal. Datasets failed preprocessing QC if more than 20% of EEG channels would require interpolation or if C3 or C4 was marked for interpolation. Linked-mastoid branches were retained only when linked mastoids (M1/M2) referencing was valid after bad-channel screening.

ICA was estimated on a 1-Hz high-pass copy of the branch-specific continuous data rather than on the ERP signal itself. The ICA copy otherwise followed the same branch-specific line-noise, lowpass, ASR, bad-channel exclusion, and reference settings up to the ICA stage. ICA decomposition used EEGLAB’s *runica* implementation of extended Infomax (Delorme & Makeig, 2004; Lee et al., 1999), with rank auditing. ICA weights were transferred to the corresponding ERP-signal data, components were classified with ICLabel, and artefact components were removed according to the branch-specific ICLabel policy. After component removal, bad channels were interpolated, the final reference was applied, and ICA fields were cleared from the final projected dataset. Full software calls, filter parameters, ICLabel thresholds, *ASR* audits, ICA-rank audits, bad-channel logs, and movement diagnostics are reported in the Supplementary Methods.

### 2.5. Epoching, quality checks, and feature extraction

Independent target-locked and response-locked epoch streams were generated from each cleaned continuous branch × participant dataset. Target-locked epochs spanned −500 to +1000 ms around target onset; at 250 Hz, the final sampled target-locked time point occurred at approximately +996 ms. Response-locked epochs spanned −800 to +500 ms around the observed response. Target-locked baseline correction followed the branch-specific cleaning-stringency bundle, using either −200 to 0 ms or −100 to 0 ms. Response-locked epochs used a −400 to −200 ms baseline in all branches. Epoch-level metadata were required for every analysed EEG dataset. Datasets were analysed only when the number of EEG epochs matched the number of corresponding metadata rows.

Epoch rejection was applied after epoching and baseline correction. Canonical branches used an absolute-amplitude threshold of 100 µV across valid EEG channels, whereas strict branches used a threshold of 75 µV. Channels marked as bad for subsequent interpolation were excluded from epoch-level artefact rejection. In gyro-sensitive branches, epochs with a gyroscope magnitude exceeding a within-dataset z-score threshold of 3 were additionally flagged. EEG analyses included only retained epochs with a valid left/right response 100-1,700 ms after target onset whose response side matched the target-defined response side.

Mean amplitudes were extracted from prespecified component windows and electrode clusters selected on the basis of established ERP literature. P1 was quantified at O1, O2, and POz between 80 and 130 ms, and posterior N1 at P7, P8, P3, P4, POz, O1, and O2 between 140 and 220 ms (Hillyard & Anllo-Vento, 1998). Fronto-central N2 was quantified at Fz, FCz, and Cz between 250 and 350 ms (Folstein & Van Petten, 2008). Centroparietal P3 was quantified at Cz, CPz, and Pz between 300 and 600 ms, and parietal P3 at P3, P4, Pz, CPz, and POz over the same interval (Polich, 2007). N2pc was computed at P7/P8 between 180 and 300 ms as contralateral-minus-ipsilateral activity relative to target location (Luck & Hillyard, 1994; Eimer, 1996; Kiss et al., 2008). Target-locked C3/C4 lateralisation was computed between 300 and 600 ms relative to response side, target-defined response side, and target location, alongside raw C4-minus-C3 asymmetry (Eimer, 1998; Praamstra & Oostenveld, 2003; Praamstra, 2007). Full component definitions are reported in the Supplementary Methods.

### 2.6. Statistical analysis and diagnostic controls

The main analysis tested whether location-response compatibility modulated target-locked EEG activity. The location-response compatibility effect was defined as the incompatible-minus-compatible contrast. Across preprocessing branches, the analysis layer was held constant: the same trial labels, component windows, electrode clusters, lateralisation definitions, inclusion rules, contrast definitions, and statistical tests were applied to every branch. Participant-level cell means were computed for each component and region of interest across combinations of cue-target interval, cue condition, flanker congruency, compatibility, and spatial-cue validity for spatial-cue trials. The compatibility effect was estimated using a strictly balanced within-participant procedure. Participants contributed to the contrast only when both compatibility conditions were available within the corresponding combination of nuisance conditions after trial-count filtering. The analysis required at least six correct retained epochs per participant-by-condition cell and balanced the incompatible-minus-compatible contrast across cue-target interval, cue condition, flanker congruency, and spatial-cue validity for spatial-cue trials.

Within each preprocessing branch, inferential tests were conducted on participant-level contrast scores using two-tailed one-sample *t*-tests against zero. Branch-level outputs included effect estimates, 95% confidence intervals, unadjusted *p* values, Holm-adjusted *p* values, and Cohen’s *d*z effect sizes. Within each branch, Holm correction was applied across the seven planned target-locked measures: P1, posterior N1, N2pc, frontocentral N2, centroparietal P3, parietal P3, and response-referenced C3/C4. Holm correction was applied separately within the secondary diagnostic families; the target-locked C3/C4 reference-frame family comprised response-side, target-defined response-side, target-location, and raw C4-minus-C3 measures. Multiverse-level summaries were descriptive and included the median effect, interquartile range, number of positive branches, number of branches with an unadjusted *p* < .05, and number of branches with Holm-adjusted *p* < .05.

The analysis hierarchy was defined before the final branch-level results were interpreted. The primary outcome was the target-locked N2pc location-response compatibility effect within the 192-branch specification space. Results across the planned component family were used to assess whether the effect was specific to the N2pc measure or was also evident in other ERP measures. C3/C4 reference-frame comparisons, response-locked analyses, pre-target controls, sign-exchange controls, trial-threshold analyses, preprocessing-decision sensitivity analyses, bootstrap uncertainty, split-half reliability, standardised measurement error, ASR-associated root-mean-square changes, movement-proxy analyses, and channel-label diagnostics were treated as interpretative, sensitivity, falsification, or data-quality analyses rather than as independent tests of the N2pc hypothesis.

## 3. Results

### 3.1. Multiverse flow, behaviour, and EEG retention

The EEG multiverse comprised 192 preprocessing branches with a constant analysis layer across branches. Reference policy was included in the specification space because reference choice was treated as a central preprocessing uncertainty source for lateralised EEG estimates. Across branches, 9,024 branch × participant preprocessing requests were launched. Of these, 8,676 generated a QC row, comprising 7,849 QC-pass rows and 827 QC-fail rows. The remaining 348 requests terminated before QC-row generation because linked-mastoid reference was requested when M1 or M2 had been marked bad in the corresponding preprocessing request; these rows are reported as pre-QC attrition rather than QC failures. All 192 branches retained analysable target-locked data. After QC filtering and restriction to the behavioural-overlap sample, 7,369 target-locked and 7,369 response-locked branch × participant datasets entered feature extraction. Analysed participants per branch ranged from 30 to 42, with a median of 39 participants per branch. The median number of correct retained target-locked epochs per branch was 10,101, IQR [9,438, 10,846], and the median decoded target-locked epoch retention rate was 95.5%. Multiverse flow, preprocessing quality-control outcomes, EEG-retention summaries, and channel-label audit results are summarised in Table 1 and reported in full in Supplementary Results.

**Table 1.** Multiverse Flow, Preprocessing Quality Checks, and EEG Retention.

| Characteristic | Value |
| --- | --- |
| Final behavioural sample | 44 participants |
| Preprocessing branches | 192 |
| Branch × participant preprocessing attempts | 9,024 |
| Preprocessing QC pass | 7,849 (87.0%) |
| Preprocessing QC fail | 827 (9.2%) |
| Pre-QC attrition | 348 (3.9%) |
| Target-locked branch × participant datasets entering feature extraction | 7,369 |
| Response-locked branch × participant datasets entering feature extraction | 7,369 |
| Analysed participants per branch | 30-42 |
| Median retained correct target-locked epochs per branch | 10,101 [IQR: 9,438-10,846] |
| Median target-locked retention rate | 95.5% [IQR: 95.3%-95.7%] |
| C3/C4 channel-label audit | Exactly one C3 and one C4 label in every analysed target- and response-locked dataset |

**Note.** Counts summarise flow through the 192-branch preprocessing multiverse after preprocessing quality checks (QC) and behavioural-overlap filtering. Percentages are relative to branch × participant preprocessing attempts where applicable. IQR = interquartile range; QC = quality checks. Target-locked and response-locked datasets refer to branch × participant datasets entering feature extraction. The C3/C4 channel-label audit confirmed exactly one C3 and one C4 label in every analysed target-locked and response-locked dataset.

Full-stream behavioural reaction-time analyses included correct responses from 100 to 1,700 ms, whereas accuracy analyses included all trials. Location-response incompatibility was not associated with either a reliable reaction-time difference (mean difference = +2.65 ms, 95% CI [−5.97, +11.27], *t*(43) = 0.62, *p* = .539), or reliable accuracy difference (mean difference = −0.76 percentage points, 95% CI [−2.08, +0.56], *t*(43) = −1.16, *p* = .254). Late responses relative to the target-locked EEG epoch boundary were rare and did not differ reliably by compatibility, mean difference = −0.25 percentage points, 95% CI [−0.78, +0.28], *t*(43) = −0.96, *p* = .341. Across branches, the median incompatible-minus-compatible EEG-retention difference was −0.39 percentage points, IQR [−0.50, −0.28]. These behavioural and retention checks did not indicate that the EEG contrast was driven by a reliable behavioural compatibility cost or by a large compatibility-related retention imbalance.

### 3.2. N2pc multiverse results

The target-locked analysis tested the location-response incompatible-minus-compatible contrast in correct retained ANT-VR epochs (Figure 2). The contrast was estimated within participant and balanced over cue-target interval, cue condition, cue-validity information where applicable, and flanker congruency.

**Figure 2.**
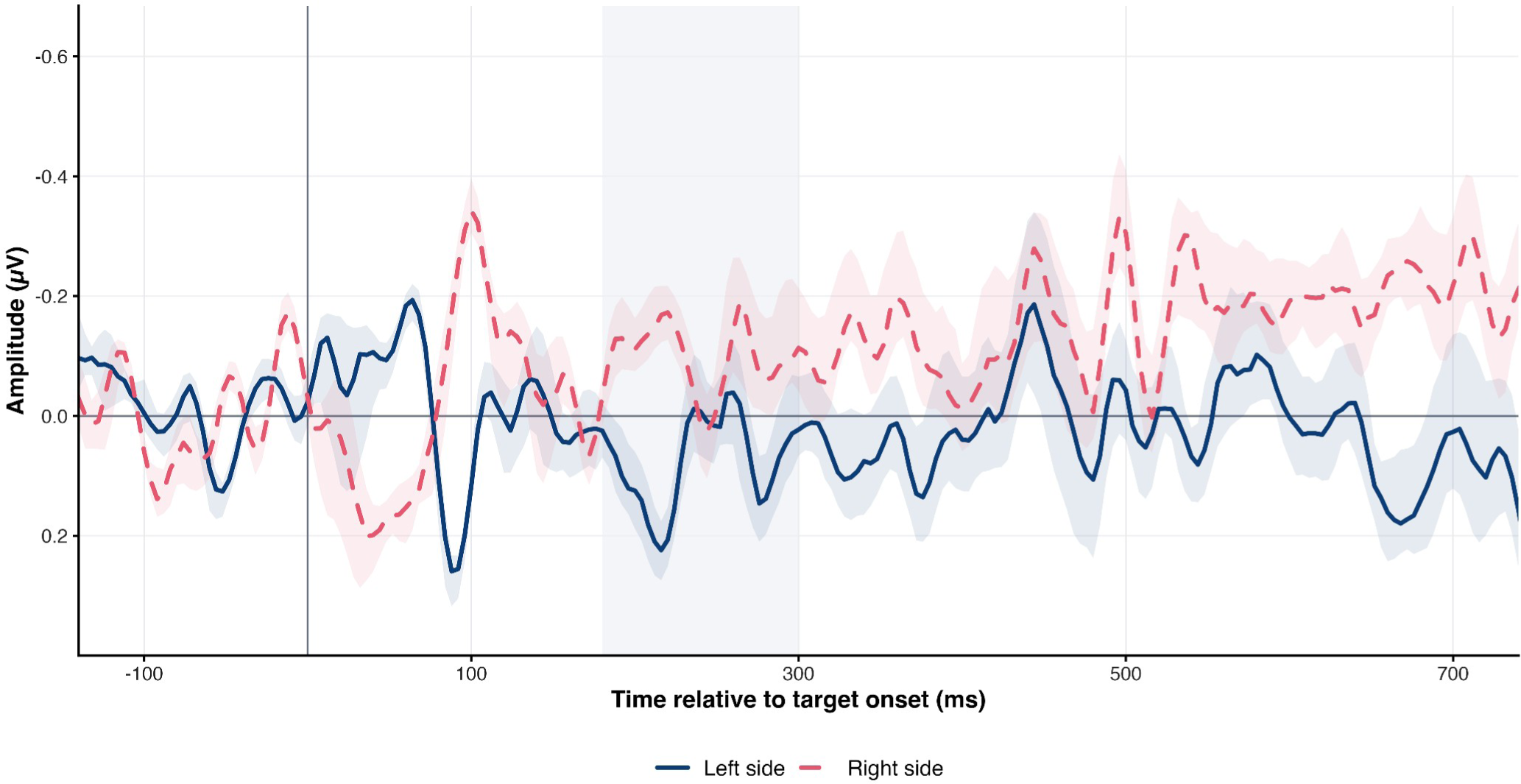
Left- and right-side posterior compatibility contrasts in the N2pc window **Note.** Lines show the median branch-level incompatible-minus-compatible waveforms separately for the left- and right-side posterior signals. Within each preprocessing branch, participant waveforms were averaged using retained trial counts as weights; the displayed lines represent the median across the 192 preprocessing specifications. The grey band marks the prespecified N2pc analysis window from 180 to 300 ms. Activity outside this window is shown descriptively and was not used for the N2pc inference. The four constituent P7 and P8 waveforms for compatible and incompatible trials are presented in Supplementary Figure S1.

The N2pc-window posterior-lateralisation contrast was estimable in all 192 preprocessing branches. Across the restricted multiverse, the N2pc incompatible-minus-compatible contrast was directionally positive in 192/192 branches, with a median effect of +0.295 µV, IQR [+0.220, +0.412], and an observed range of approximately +0.05 to +0.79 µV. Participant-resampling bootstrap uncertainty placed the median branch effect above zero, 95% CI [+0.020, +0.594] µV. The N2pc-window posterior-lateralisation contrast met the nominal *p* < .05 criterion in 79/192 branches and survived within-branch Holm correction for the planned component family in 42/192 branches. The N2pc specification curve is shown in Figure 3; component-level robustness across the planned target-locked family is reported in Table 2.

**Table 2.** Component-Level Target-Locked Multiverse Robustness.

| Measure | Median effect [IQR] | Positive branches | Holm p < .05 branches |
| --- | --- | --- | --- |
| P1 | 0.08 μV [0.03-0.15] | 181/192 | 0/192 |
| Posterior N1 | 0.06 μV [0.02-0.11] | 165/192 | 0/192 |
| N2pc | 0.30 μV [0.22-0.41] | 192/192 | 42/192 |
| Frontocentral N2 | -0.14 μV [-0.25 to -0.07] | 5/192 | 0/192 |
| Centroparietal P3 | 0.01 μV [-0.14 to 0.10] | 100/192 | 0/192 |
| Parietal P3 | 0.00 μV [-0.10 to 0.11] | 91/192 | 0/192 |
| Response-referenced C3/C4 | 0.23 μV [0.02-0.47] | 157/192 | 0/192 |

**Note.** Effects are branch-level incompatible-minus-compatible estimates across the 192 preprocessing branches. Effects are reported in microvolts. Positive branches indicate the number of branches in which the incompatible-minus-compatible estimate was greater than zero. Holm p < .05 branches indicate the number of branches surviving Holm correction within the planned component family. Branch counts are descriptive robustness indicators across dependent preprocessing specifications and should not be interpreted as independent replications. IQR = interquartile range; N1 = posterior N1; N2 = frontocentral N2; N2pc = N2 posterior contralateral component; P1 = posterior P1; P3 = P3 event-related potential component.

**Figure 3.**
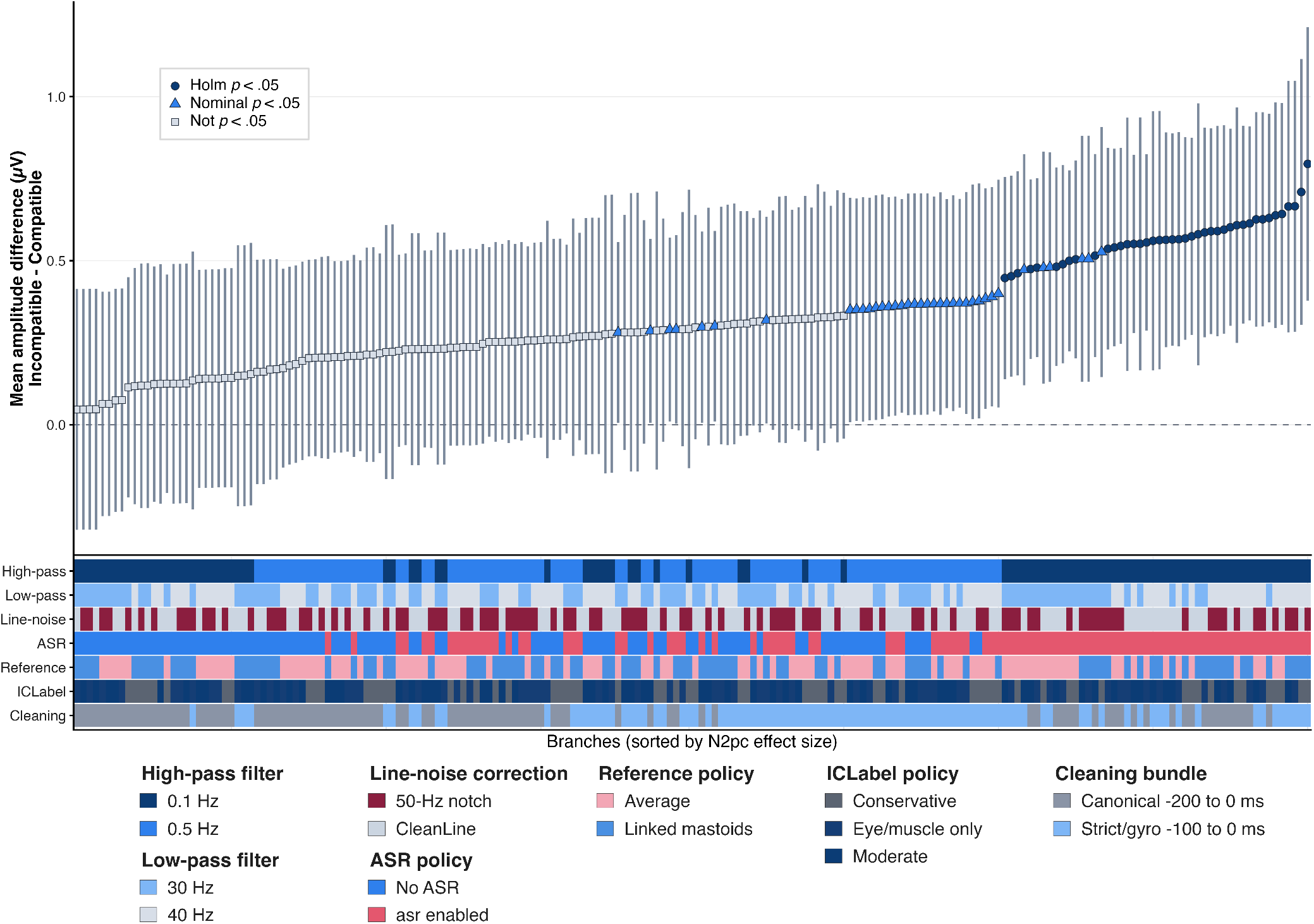
N2pc Specification Curve with Preprocessing-Decision Heatmap **Note.** The upper panel shows branch-level estimates of the N2pc-window posterior-lateralisation compatibility effect, defined as the incompatible-minus-compatible contrast. Branches are ordered from the smallest to the largest estimated effect. Points represent preprocessing specifications, error bars indicate 95% confidence intervals, and point symbols distinguish branches with Holm-adjusted p < .05, nominal p < .05, and p ≥ .05. The dashed horizontal line denotes a null effect of zero. The lower panel shows the preprocessing decisions defining each branch, with each column aligned with the corresponding estimate in the upper panel. The categories represented in the heatmap are identified in the key below the figure. All branches comprise alternative preprocessing specifications applied to the same participants and data and should therefore not be interpreted as independent replications.

Because branches reused the same participants, raw data, task labels, feature definitions, and statistical tests, these counts are descriptive robustness summaries rather than independent replications. Thus, the N2pc-window posterior-lateralisation contrast showed high directional stability across a bounded set of dependent, defensible preprocessing specifications, rather than indicating that the effect replicated 192 times.

### 3.3. Component-family specificity

The incompatible-minus-compatible contrast was also estimated for the planned target-locked component family: P1, posterior N1, N2pc, frontocentral N2, centroparietal P3, parietal P3, and response-referenced C3/C4. These analyses tested whether the N2pc effect was broadly distributed across ERP components or concentrated in a theoretically relevant lateralised target-selection measure.

N2pc was the only component-family target with Holm-corrected support in a subset of branches. P1 and posterior N1 showed small positive median effects but no Holm-significant branches. Frontocentral N2 showed a small negative median effect and no Holm-significant branches. Centroparietal and parietal P3 estimates were centred close to zero. The response-referenced C3/C4 analysis showed a median effect of +0.231 µV, IQR [+0.020, +0.467], was positive in 157/192 branches, and survived Holm correction in 0/192 branches. Thus, within the planned component-family analysis, the strongest branch-stable evidence was concentrated in the N2pc-window posterior-lateralisation measure rather than distributed across early sensory, frontocentral conflict-related, P3-family, or response-referenced C3/C4 measures. These component-family contrasts are summarised numerically in Table 2 and visualised in Figure 4.

**Figure 4.**
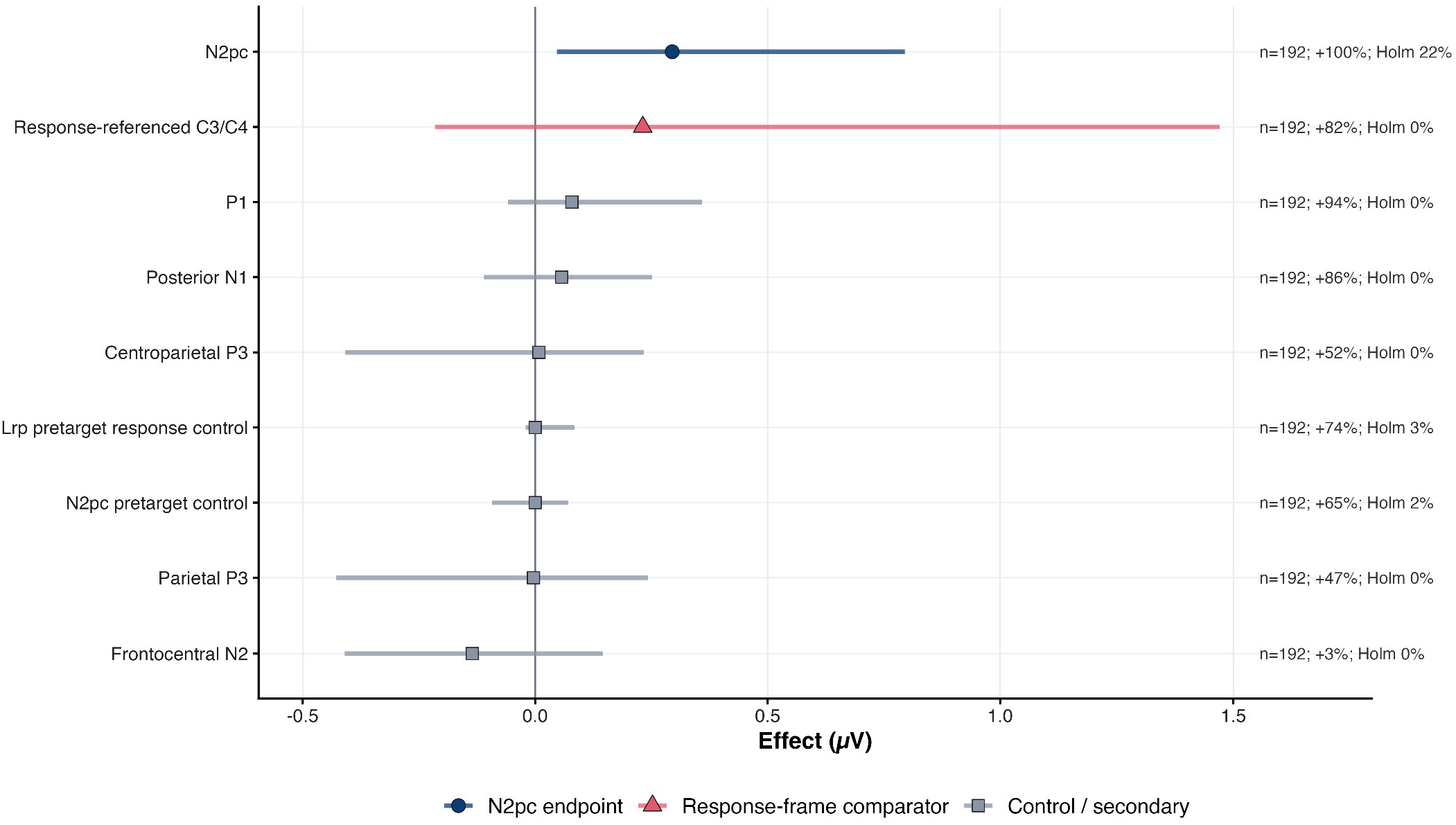
Component- and Target-family Comparison **Note.** Points show median branch effects for the planned target-locked component family together with selected pre-target control measures in the incompatible-minus-compatible contrast. Horizontal intervals show the exported branch range. Annotations summarise the proportion of branches in the positive direction and the proportion surviving Holm correction. Branch-level summaries are descriptive because preprocessing specifications share the same participants and raw data.

### 3.4. C3/C4 reference-frame decomposition

C3/C4 decomposition analyses were used as sensor-level reference-frame analyses. They were not treated as source-localising evidence, proof of an isolated motor generator, or replacements for the N2pc endpoint. Instead, they tested whether C3/C4 compatibility effects were better described in response-side, target-defined response-side, target-location, or raw hemispheric coordinates.

Target-location-referenced C3/C4 showed the strongest decomposition effect, with a median incompatible-minus-compatible estimate of +0.528 µV, IQR [+0.45, +0.65]. The effect was positive in 192/192 branches and survived Holm correction within the four-measure target-locked C3/C4 reference-frame family in 108/192 branches. By contrast, response-referenced C3/C4 showed a weaker median effect of +0.231 µV, IQR [+0.020, +0.467], was positive in 157/192 branches, and survived the same reference-frame-family Holm correction in 0/192 branches. Target-defined response-side C3/C4 was identical to response-referenced C3/C4 because the feature set used here was restricted to correct retained trials, for which observed response side and target-defined response side were structurally coupled. Raw C4-minus-C3 asymmetry was centred near zero, median = +0.011 µV, IQR [−0.09, +0.06], with 102/192 positive branches and no Holm-significant branches. The target-locked reference-frame decomposition results are summarised in Table 3 and visualised in Figure 5. While the target-location-referenced C3/C4 effect was stronger and more consistently corrected across branches than the N2pc-window posterior-lateralisation contrast, it was retained as a secondary reference-frame analysis because C3/C4 lateralisation is less anatomically and functionally specific in the present 24-channel mobile EEG montage.

**Table 3.**
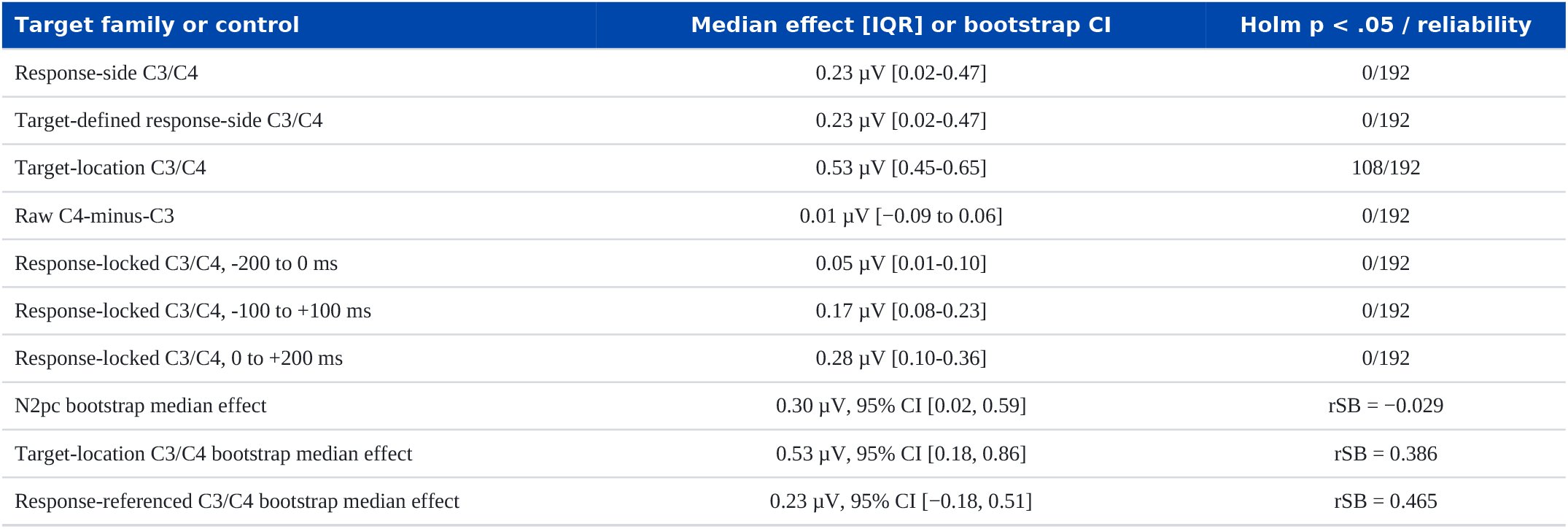
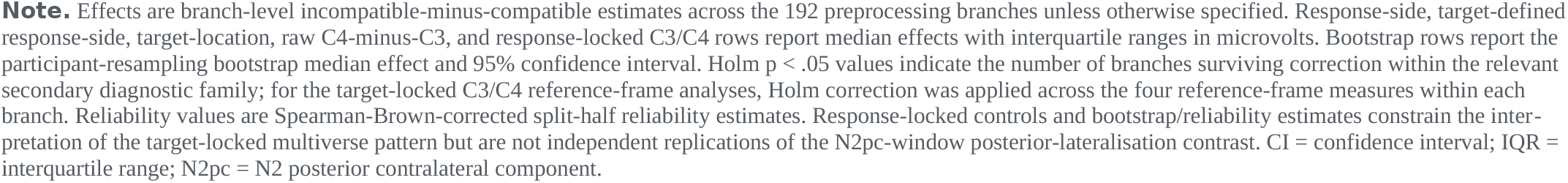
Lateralised Reference-Frame and Control Analyses.

| Target family or control | Median effect [IQR] or bootstrap CI | Holm p < .05 / reliability |
| --- | --- | --- |
| Response-side C3/C4 | 0.23 μV [0.02-0.47] | 0/192 |
| Target-defined response-side C3/C4 | 0.23 μV [0.02-0.47] | 0/192 |
| Target-location C3/C4 | 0.53 μV [0.45-0.65] | 108/192 |
| Raw C4-minus-C3 | 0.01 μV [-0.09 to 0.06] | 0/192 |
| Response-locked C3/C4, -200 to 0 ms | 0.05 μV [0.01-0.10] | 0/192 |
| Response-locked C3/C4, -100 to +100 ms | 0.17 μV [0.08-0.23] | 0/192 |
| Response-locked C3/C4, 0 to +200 ms | 0.28 μV [0.10-0.36] | 0/192 |
| N2pc bootstrap median effect | 0.30 μV, 95% CI [0.02, 0.59] | rSB = -0.029 |
| Target-location C3/C4 bootstrap median effect | 0.53 μV, 95% CI [0.18, 0.86] | rSB = 0.386 |
| Response-referenced C3/C4 bootstrap median effect | 0.23 μV, 95% CI [-0.18, 0.51] | rSB = 0.465 |

**Note.** Effects are branch-level incompatible-minus-compatible estimates across the 192 preprocessing branches unless otherwise specified. Response-side, target-defined response-side, target-location, raw C4-minus-C3, and response-locked C3/C4 rows report median effects with interquartile ranges in microvolts. Bootstrap rows report the participant-resampling bootstrap median effect and 95% confidence interval. Holm $p < .05$ values indicate the number of branches surviving correction within the relevant secondary diagnostic family; for the target-locked C3/C4 reference-frame analyses, Holm correction was applied across the four reference-frame measures within each branch. Reliability values are Spearman-Brown-corrected split-half reliability estimates. Response-locked controls and bootstrap/reliability estimates constrain the interpretation of the target-locked multiverse pattern but are not independent replications of the N2pc-window posterior-lateralisation contrast. CI = confidence interval; IQR = interquartile range; N2pc = N2 posterior contralateral component.

**Figure 5.**
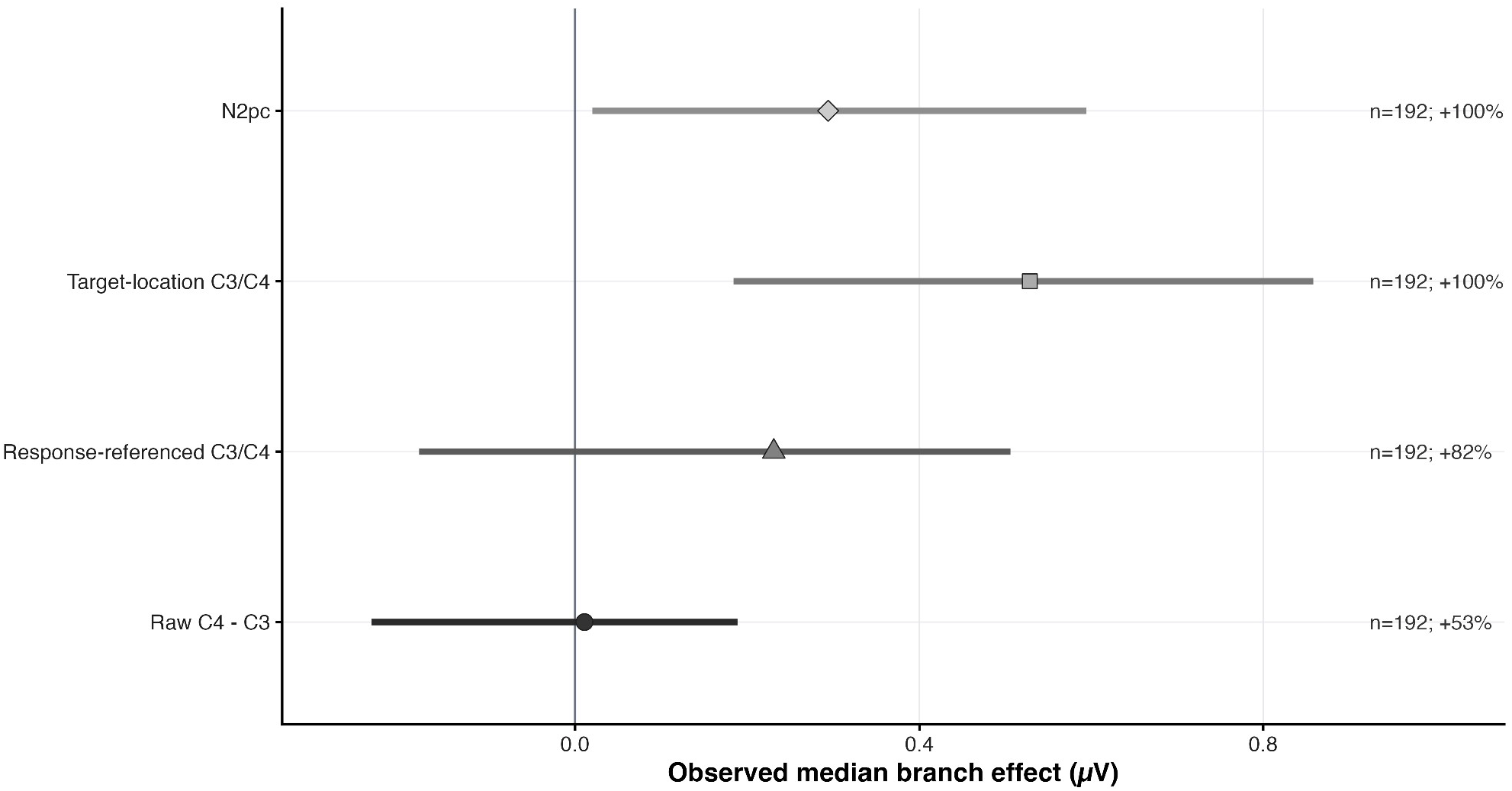
Comparison of Lateralisation Reference Frames **Note.** Points show observed median branch effects for N2pc, target-location-referenced C3/C4, response-referenced C3/C4, and raw C4-minus-C3 asymmetry. Horizontal intervals indicate 95% participant-bootstrap intervals. The figure evaluates whether the compatibility effect is expressed similarly across lateralisation reference frames.

The decomposition therefore argues against treating the compatibility effect as a simple response-preparation effect. The stronger target-location-referenced C3/C4 pattern indicates that the lateralised central signal was reference-frame sensitive and more strongly constrained by target location than by response side alone. Together with the N2pc result, this pattern is consistent with target-locked spatial-selection or target-location-linked lateralisation, while not isolating a single spatial, premotor, or motor source.

### 3.5. Diagnostic controls, uncertainty, and sensitivity

Response-locked and pre-target analyses tested whether the target-locked pattern could be explained by response-related lateralisation or pre-stimulus asymmetry. Response-locked C3/C4 effects were positive in windows before, around, and after response onset but did not survive Holm correction in any branch. Pre-target negative-control analyses in the −200 to 0 ms target-locked window were centred close to zero for N2pc, response-referenced C3/C4, target-location-referenced C3/C4, and raw C4-minus-C3, and did not reproduce the target-locked pattern. Participant-level sign-exchange diagnostics supported the observed N2pc and target-location-referenced C3/C4 median effects relative to the sign-exchange null (*p* = .034 and *p* = .010, respectively), but not the corresponding directional-stability statistics (*p* = .180 and *p* = .265, respectively), whereas raw C4-minus-C3 did not show comparable evidence. These control analyses are reported in full in the Supplementary Results; key response-locked, bootstrap, and reliability summaries are included in Table 3.

Measurement-quality and artefact diagnostics were used to contextualise the multiverse results. For the N2pc measure, the median branch-level standardised measurement error (SME) was 0.278 µV, IQR [0.258, 0.309]. Across branch × participant observations, N2pc magnitude was only weakly associated with mean SME, *r* = .060, and was not materially associated with the prespecified C3/C4 movement proxy, *r* = .027. The movement-proxy definition and detailed artefact subspace reconstruction diagnostics are reported in the Supplementary Methods and Results.

Bootstrap and reliability analyses further constrained the interpretation. The median N2pc branch effect was 0.295 µV, with a 95% bootstrap CI of [0.020, 0.594]. Target-location-referenced C3/C4 also had a positive median branch effect of 0.528 µV, 95% bootstrap CI [0.184, 0.858]. In contrast, the bootstrap intervals for response-referenced C3/C4 and raw C4-minus-C3 included zero. Repeated stratified split-half reliability was weak for N2pc, with a median split-half *r* of −.01 and a Spearman-Brown-corrected reliability of −.03. The N2pc result should therefore be interpreted as a group-level experimental effect within the restricted preprocessing multiverse, rather than as an individual-difference measure or biomarker candidate.

Preprocessing-decision and trial-threshold sensitivity analyses were consistent with this interpretation. Descriptively, the largest differences in the median N2pc estimate were associated with ASR policy, the cleaning-stringency bundle, and the high-pass filter, whereas low-pass filter, line-noise attenuation, reference policy, and ICLabel policy showed smaller differences. Trial-threshold analyses repeated the target-family estimates using minimum cell thresholds of 1, 3, 6, 8, and 10 trials. The median N2pc-window posterior-lateralisation contrast remained positive across thresholds, although the proportion of branches with an unadjusted *p* < .05 varied. Target-location-referenced C3/C4 showed greater directional stability across thresholds, whereas response-referenced C3/C4 was more sensitive to the trial-count threshold. Overall, the multiverse results support a group-level target-locked lateralised location-response compatibility effect, most consistently observed for N2pc and target-location-referenced C3/C4, while indicating that effect magnitude and inferential strength remained partly sensitive to preprocessing and trial-count decisions.

## 4. Discussion

### 4.1. Principal finding

The present study examined the neural processes associated with location-response compatibility in an immersive object-based implementation of the ANT-R. Across a restricted preprocessing multiverse used to assess robustness to alternative defensible preprocessing choices, the posterior-lateralisation contrast in the N2pc time window showed high directional stability, with Holm-corrected support in a subset of specifications. Because N2pc was scored as contralateral-minus-ipsilateral activity relative to target location, positive incompatible-minus-compatible values indicate a shift of the lateralised difference towards positivity in incompatible trials. The result should therefore not be interpreted as an enhanced conventional negative-going N2pc. Instead, location-response compatibility modulated target-locked posterior lateralisation during the N2pc time window.

This result extends previous electrophysiological research using the ANT. Earlier ANT studies with conventional two-dimensional stimuli have linked alerting and orienting to early target-related P1/N1 modulation, and executive conflict or inhibition to later P3-related activity (Neuhaus et al., 2010; Galvao-Carmona et al., 2014; Santhana Gopalan et al., 2019). The present study focused on a different contrast and identified the most consistent compatibility-related effect in a lateralised posterior measure rather than in the early sensory, frontocentral N2, or P3 measures examined. Thus, the current findings do not reproduce a previously established ANT N2pc compatibility effect; rather, they extend ANT electrophysiology by showing that location-response compatibility can modulate posterior lateralisation when the task is implemented with meaningful objects in immersive VR.

The interpretation of this effect is consistent with established accounts of N2pc as a marker of spatially selective processing of lateralised target objects (Luck & Hillyard, 1994; Eimer, 1996; Kiss et al., 2008). Together with the stronger target-location-referenced C3/C4 effect, the pattern suggests that compatibility influenced lateralised activity associated with target selection or target-location coding. However, because the effect represented a positive shift in the incompatible-minus-compatible contrast rather than a larger conventional negative-going N2pc, the data do not establish enhanced attentional selection in incompatible trials. They show more cautiously that compatibility altered posterior lateralisation within the temporal interval typically associated with spatially selective target processing.

Although reaction time and accuracy did not show reliable compatibility differences, this dissociation is consistent with the broader rationale for using ERPs in ANT research: electrophysiological measures can distinguish processing stages that are combined at the behavioural endpoint (Neuhaus et al., 2010; Galvao-Carmona et al., 2014). The current result therefore indicates a group-level neural modulation in the absence of a reliable behavioural compatibility cost, rather than a covert behavioural effect inferred solely from non-significance.

### 4.2. Spatial-selection lateralisation rather than a simple response-preparation account

The C3/C4 reference-frame comparisons constrained the interpretation of the lateralised effects. These analyses tested whether compatibility-related C3/C4 modulation was better described relative to observed response side, target-defined response side, target location, or raw hemispheric coordinates. Across the multiverse, the evidence did not support a specifically response-referenced account. Response-referenced C3/C4 was weaker than target-location-referenced C3/C4 and did not show Holm-corrected branch-level support. The stronger target-location-referenced effect therefore argues against interpreting the compatibility effect solely as a conventional lateralised-readiness-potential-like response-preparation effect.

This conclusion is relevant to previous electrophysiological work on spatial compatibility. In an arrow-based Simon task, Cespón et al. (2012) found that compatibility modulated the lateralised readiness potential and N2cc but not N2pc, consistent with a substantial contribution of response selection and response monitoring. The present profile differed: the response-referenced C3/C4 effect was comparatively weak, whereas posterior N2pc-window and target-location-referenced C3/C4 effects were more consistent. This difference should not be taken as evidence that motor processing was absent. Instead, it suggests that the neural expression of compatibility in ANT-VR was not restricted to the response-centred profile reported in a conventional arrow-based Simon task.

At the same time, target-location-referenced C3/C4 should not be treated as process-pure evidence of posterior spatial attention. Lateralised central activity in spatial compatibility tasks can reflect overlapping visuospatial, premotor, response-selection, and sensorimotor contributions (Eimer, 1998; Praamstra & Oostenveld, 2003; Praamstra, 2007). Moreover, target identity and required response side were structurally coupled on correct trials. The most defensible conclusion is therefore that the central lateralised signal was more strongly organised by target location than by response side alone. Together with the posterior result, this supports a target-location-linked interpretation while preserving uncertainty regarding the relative contributions of posterior selection, premotor coding, and motor preparation.

### 4.3. Component specificity and theoretical interpretation

The comparison across ERP measures distinguished among the candidate accounts outlined in the Introduction. P1 and posterior N1 effects would have been consistent with early sensory-spatial modulation, whereas frontocentral N2 and P3 effects would have been more consistent with cognitive-control or mismatch processing and stimulus-evaluation or context-updating processes, respectively (Hillyard & Anllo-Vento, 1998; Folstein & Van Petten, 2008; Polich, 2007). The multiverse did not place its strongest evidence in these component families. Instead, the most consistent effects occurred in posterior lateralisation during the N2pc window and in target-location-referenced C3/C4.

The N2pc literature provides an important context for this finding. N2pc is commonly used as an index of spatially selective processing at task-relevant lateralised locations (Eimer, 1996; Kiss et al., 2008), but the selected representation need not be an abstract position in otherwise empty space. Woodman et al. (2009) observed anticipatory N2pc activity when cued locations were marked by placeholder objects, but not when the corresponding spatial locations lacked object structure. Eimer and Grubert (2014) further showed that N2pc activity can distinguish temporally evolving feature-based and object-based stages of selection. These findings support the plausibility that phones and tablets embedded in the ANT-VR scene were selected as spatially situated objects rather than represented solely as left-right coordinates.

This literature does not, however, establish that the present compatibility modulation was specifically object-based. The ANT-VR design did not independently manipulate object structure and spatial position, and target identity, target location, and response mapping were intertwined. Moreover, the current effect did not take the form of a conventional enhancement of the negative-going N2pc. The findings therefore support the narrower conclusion that compatibility influenced posterior lateralisation associated with selection or coding of a lateralised object. They do not demonstrate a process-pure object-based attention mechanism.

The findings are also compatible with spatial stimulus-response compatibility accounts in the broader sense that a task-irrelevant spatial attribute acquired functional significance because it overlapped with a task-defined response code (Kornblum et al., 1990; Lu & Proctor, 1995). However, ANT-VR used an immersive object-based implementation, and the contrast with the response-centred Simon profile discussed above could reflect stimulus format, object structure, perceptual-action context, analysis definitions, or combinations of these factors. The present design cannot isolate which difference was responsible.

### 4.4. Multiverse robustness and preprocessing dependence

A key contribution of the present study is that the principal interpretation was evaluated across a restricted preprocessing multiverse rather than through a single fixed pipeline. This was particularly relevant because filtering, referencing, artefact correction, baseline adjustment, and the number of retained trials can influence ERP estimates and their precision (Clayson et al., 2021), while the effectiveness of artefact removal can depend on the selected rejection or correction parameters (Chang et al., 2020). Accordingly, the multiverse tested whether the interpretation remained stable across defensible choices in filtering, line-noise attenuation, artefact correction, referencing, ICA-based component rejection, and cleaning stringency. Because the branches reused the same participants, recordings, task labels, component definitions, and statistical tests, they were treated as dependent specifications rather than independent replications.

The resulting evidence was both robust and qualified. Posterior lateralisation during the N2pc window was positive in all 192 branches, and participant resampling placed the median branch effect above zero. However, only a subset of branches survived Holm correction, and the magnitude and inferential strength of the effect varied across preprocessing decisions. The appropriate conclusion is therefore not that the result replicated 192 times, but that its direction was stable across many defensible preprocessing specifications.

This distinction also limits the strength of the theoretical inference. The multiverse reduces concern that the observed pattern depended entirely on one preferred preprocessing pipeline. Yet it does not remove uncertainty about effect magnitude or component identity.

### 4.5. Implications for ANT-VR and object-based attention

The present findings extend previous work showing that the major behavioural structure of the ANT-R can be preserved when the task is translated into immersive VR (Tekampe et al., 2023, 2026). Previous ANT ERP studies have reported attention-related N1, N2, and P3 effects with conventional arrow or fish stimuli (Neuhaus et al., 2010; Abundis-Gutiérrez et al., 2014; Galvao-Carmona et al., 2014; Santhana Gopalan et al., 2019), whereas N2pc research has demonstrated spatially selective processing of targets defined as shapes, feature conjunctions, and perceptual objects (Woodman et al., 2009; Eimer & Grubert, 2014). The present study connects these literatures by showing that a target-locked lateralised EEG effect remains observable when the ANT-R is implemented with familiar objects in an immersive scene. This represents generalisation across task implementation rather than direct replication of an identical component contrast.

Phones and tablets were not arbitrary arrows or colour patches; they were recognisable objects positioned within a virtual apartment. Their physical locations may therefore have affected how the target was selected or spatially coded, even though participants responded using controllers rather than directly manipulating the objects (Woodman et al., 2009; Eimer & Grubert, 2014). A recent VR Simon study found larger behavioural compatibility effects for reachable than unreachable stimuli, indicating that task-irrelevant spatial correspondence can depend on the potential for interaction with objects in three-dimensional space (Paavola et al., 2026). Although the present task did not manipulate reachability and did not produce a reliable behavioural compatibility effect, that finding reinforces the broader possibility that immersive perceptual-action context alters how spatial compatibility is expressed.

The current data do not permit a strong affordance interpretation. Object identity, spatial position, and response mapping were not independently manipulated, and participants responded via controller button presses rather than acting directly on the virtual objects. The findings nevertheless suggest that translating an established attention paradigm into VR need not eliminate its time-resolved neural signatures. Instead, the translation may preserve lateralised selection-related processing while changing whether and how that processing appears in behavioural performance and response-referenced neural measures.

### 4.6. Limitations

Several limitations constrain the interpretation. Preserving the mobile ANT-VR recording context introduced methodological challenges that are less prominent in conventional stationary EEG paradigms. Interpretation may be complicated by controller and headset movement, cap displacement, muscle activity, marker-timing uncertainty, and response-related artefacts. The constituent contralateral and ipsilateral waveforms, pre-target controls, response-locked controls, movement diagnostics, and preprocessing-sensitivity analyses are therefore important for establishing that the posterior effect cannot be attributed straightforwardly to a single response-related or preprocessing-specific explanation.

The findings describe a neural profile within the specific perceptual-action context of ANT-VR and should not be interpreted as evidence for a general VR-induced change in attention mechanisms. In addition, the task did not independently manipulate object status, physical location, target identity, and response side. The results can therefore be interpreted in relation to object-based and spatial-selection research, but they cannot establish that object-based selection caused the observed modulation. The absence of a neutral centrally located compatibility condition further limits direct comparison with Simon studies that used neutral-position subtraction to remove common motor activity and isolate posterior N2pc and central N2cc from overlapping LRP-related activity (Cespón et al., 2012).

The multiverse also places specific constraints on inference. Because preprocessing branches shared participants and raw recordings, branch counts index robustness across specifications rather than replication frequency. In addition, split-half reliability was weak at the individual level. The findings therefore support a group-level experimental effect but not an individual-difference or biomarker interpretation.

Further limitations concern spatial specificity and generalisability. Given the limited spatial specificity of surface-recorded ERPs, the 24-channel montage constrains source-level separation of overlapping posterior, central, premotor, and motor activity (Hillyard & Anllo-Vento, 1998; Praamstra & Oostenveld, 2003). The sample also consisted of young university participants, limiting generalisability to older, clinical, or more heterogeneous populations. These limitations do not negate the observed posterior-lateralisation effect, but they define its scope: a group-level location-response compatibility modulation in an immersive object-based ANT-R implementation, expressed most consistently in target-locked lateralised measures.

### 4.7. Conclusion

Location-response compatibility in immersive ANT-VR modulated target-locked lateralised EEG activity. Across the restricted preprocessing multiverse, posterior lateralisation during the N2pc window showed high directional stability, and target-location-referenced C3/C4 provided convergent reference-frame evidence. Response-referenced C3/C4 was weaker, arguing against a solely response-preparation-centred interpretation.

The findings extend previous ANT electrophysiology and N2pc object-selection research to an immersive object-based task. They do not demonstrate an enhanced conventional N2pc or directly replicate an established N2pc Simon effect. Instead, they support a bounded account in which location-response compatibility can alter lateralised target-selection- or target-location-related processing even when behavioural compatibility differences are weak or absent.

## 5. Statements and Declarations

### Funding

This work was supported by the University Fund Limburg/SWOL (grant CoBes.23.019 to L.H. and M.S.), with additional institutional support from Maastricht University during data collection and the University of Luxembourg during data analysis and manuscript preparation.

### Competing Interests

The authors declare that they have no relevant financial or non-financial interests to disclose.

### Ethics approval

This study was performed in accordance with the ethical standards laid down in the 1964 Declaration of Helsinki and its later amendments. Ethical approval was granted by the Ethics Review Committee Psychology and Neuroscience at Maastricht University (249_29_02_2022).

### Consent to participate

Written informed consent was obtained from all participants prior to their inclusion in the study.

### Data Availability

The de-identified data supporting the findings of this study will be made publicly available upon publication in an EU-based repository, in accordance with GDPR and the approved ethics protocol.

### Open Practices Statement

The study was not preregistered. The de-identified data supporting the findings of this study, experimental task application, preprocessing and analysis pipeline, figure generation pipeline, together with the source-data files used to generate the figures will be made available upon publication.

### AI-assisted tools

OpenAI tools were used only for outline, language refinement and improvement of readability. The authors reviewed and verified all output and take responsibility for the final manuscript.

## Acknowledgements

This article is dedicated to the author’s cats and owl, with gratitude for their limitless love and companionship.

## 6. Abbreviations

ANT-R: Attention Network Test-Revised
ANT-VR: virtual-reality adaptation of the Attention Network Test-Revised
ASR: artefact subspace reconstruction
CI: confidence interval
EEG: electroencephalography
ERP: event-related potential
FDR: false discovery rate
ICA: independent component analysis
IQR: interquartile range
LRP: lateralised readiness potential
N1: posterior N1 event-related potential component
N2: frontocentral N2 eventrelated potential component
N2pc: N2 posterior contralateral component
P1: posterior P1 event-related potential component
P3: P3 event-related potential component
PCA: principal component analysis
QC: quality check
RMS: root mean square
ROI: region of interest
SME: standardised measurement error
VR: virtual reality
XDF: Extensible Data Format

---

| Author | Author Contributions |
| --- | --- |
| **D. L. Tekampe** | Conceptualization, Investigation, Project administration, Data curation, Formal analysis, Methodology, Software, Visualization, Writing - original draft, Writing - review & editing. |
| **P. S. Santangelo** | Supervision, Validation, Writing - review & editing. |
| **A. Sulaj** | Investigation, Project administration, Writing - review & editing. |
| **P. Tekampe** | Software, Writing - review & editing. |
| **M. Schwartze** | Funding acquisition, Investigation, Project administration, Resources, Supervision, Validation, Writing - review & editing. |
| **L. Hausfeld** | Funding acquisition, Investigation, Project administration, Resources, Supervision, Validation, Writing - review & editing. |

## Notes

### Competing Interest Statement

The authors have declared no competing interest.

### Summary of Updates

This revision was submitted solely to correct the authors institutional affiliations. These changes are administrative in nature and do not affect the scientific content, analyses, results, or conclusions of the preprint.

## References

Abundis-Gutiérrez, A., Checa, P., Castellanos, C., & Rueda, M. R. (2014). Electrophysiological correlates of attention networks in childhood and early adulthood. Neuropsychologia, 57, 78–92. 10.1016/j.neuropsychologia.2014.02.013

Bigdely-Shamlo, N., Mullen, T., Kothe, C., Su, K.-M., & Robbins, K. A. (2015). The PREP pipeline: Standardized preprocessing for large-scale EEG analysis. Frontiers in Neuroinformatics, 9, Article 16. 10.3389/fninf.2015.00016

Bohil, C. J., Alicea, B., & Biocca, F. A. (2011). Virtual reality in neuroscience research and therapy. Nature Reviews Neuroscience, 12(12), 752–762. 10.1038/nrn3122

Cespón, J., Galdo-Álvarez, S., & Díaz, F. (2012). The Simon effect modulates N2cc and LRP but not the N2pc component. International Journal of Psychophysiology, 84(2), 120–129. 10.1016/j.ijpsycho.2012.01.019

Chang, C.-Y., Hsu, S.-H., Pion-Tonachini, L., & Jung, T.-P. (2020). Evaluation of artifact subspace reconstruction for automatic artifact components removal in multi-channel EEG recordings. IEEE Transactions on Biomedical Engineering, 67(4), 1114–1121. 10.1109/TBME.2019.2930186

Clayson, P. E., Baldwin, S. A., Rocha, H. A., & Larson, M. J. (2021). The data-processing multiverse of event-related potentials (ERPs): A roadmap for the optimization and standardization of ERP processing and reduction pipelines. NeuroImage, 245, 118712. 10.1016/j.neuroimage.2021.118712

Delorme, A., & Makeig, S. (2004). EEGLAB: an open source toolbox for analysis of single-trial EEG dynamics including independent component analysis. Journal of Neuroscience Methods, 134(1), 9–21. 10.1016/j.jneumeth.2003.10.009

Eimer, M. (1996). The N2pc component as an indicator of attentional selectivity. Electroencephalography and Clinical Neurophysiology, 99(3), 225–234. 10.1016/0013-4694(96)95711-9

Eimer, M. (1998). The lateralized readiness potential as an on-line measure of central response activation processes. Behavior Research Methods, Instruments, & Computers, 30(1), 146–156. 10.3758/BF03209424

Eimer, M., & Grubert, A. (2014). The gradual emergence of spatially selective target processing in visual search: From feature-specific to object-based attentional control. Journal of Experimental Psychology: Human Perception and Performance, 40(5), 1819–1831. 10.1037/a0037387

Fan, J., Gu, X., Guise, K. G., Liu, X., Fossella, J., Wang, H., & Posner, M. I. (2009). Testing the behavioral interaction and integration of attentional networks. Brain and Cognition, 70(2), 209–220. 10.1016/j.bandc.2009.02.002

Fan, J., McCandliss, B. D., Sommer, T., Raz, A., & Posner, M. I. (2002). Testing the efficiency and independence of attentional networks. Journal of Cognitive Neuroscience, 14(3), 340–347. 10.1162/089892902317361886

Folstein, J. R., & Van Petten, C. (2008). Influence of cognitive control and mismatch on the N2 component of the ERP: A review. Psychophysiology, 45(1), 152–170. 10.1111/j.1469-8986.2007.00602.x

Galvao-Carmona, A., González-Rosa, J. J., Hidalgo-Muñoz, A. R., Páramo, D., Benítez, M. L., Izquierdo, G., & Vázquez-Marrufo, M. (2014). Disentangling the attention network test: Behavioral, event related potentials, and neural source analyses. Frontiers in Human Neuroscience, 8, Article 813. 10.3389/fnhum.2014.00813

Hillyard, S. A., & Anllo-Vento, L. (1998). Event-related brain potentials in the study of visual selective attention. Proceedings of the National Academy of Sciences of the United States of America, 95(3), 781–787. 10.1073/pnas.95.3.781

Kilteni, K., Groten, R., & Slater, M. (2012). The sense of embodiment in virtual reality. Presence: Teleoperators and Virtual Environments, 21(4), 373–387. 10.1162/PRES_a_00124

Kiss, M., van Velzen, J., & Eimer, M. (2008). The N2pc component and its links to attention shifts and spatially selective visual processing. Psychophysiology, 45(2), 240–249. 10.1111/j.1469-8986.2007.00611.x

Kothe, C., Shirazi, S. Y., Stenner, T., Medine, D., Boulay, C., Grivich, M. I., Artoni, F., Mullen, T., Delorme, A., & Makeig, S. (2025). The lab streaming layer for synchronized multimodal recording. Imaging Neuroscience (Cambridge, Mass.), 3, IMAG.a.136. 10.1162/IMAG.a.136

Kornblum, S., Hasbroucq, T., & Osman, A. (1990). Dimensional overlap: Cognitive basis for stimulus-response compatibility—A model and taxonomy. Psychological Review, 97(2), 253–270. 10.1037/0033-295X.97.2.253

Lee, T. W., Girolami, M., & Sejnowski, T. J. (1999). Independent component analysis using an extended infomax algorithm for mixed subgaussian and supergaussian sources. Neural Computation, 11(2), 417–441. 10.1162/089976699300016719

Lu, C.-H., & Proctor, R. W. (1995). The influence of irrelevant location information on performance: A review of the Simon and spatial Stroop effects. Psychonomic Bulletin & Review, 2(2), 174–207. 10.3758/BF03210959

Luck, S. J., & Hillyard, S. A. (1994). Electrophysiological correlates of feature analysis during visual search. Psychophysiology, 31(3), 291–308. 10.1111/j.1469-8986.1994.tb02218.x

Mullen, T. R. (2012). CleanLine [Computer software]. Neuroimaging Tools and Re-sources Collaboratory. https://www.nitrc.org/projects/cleanline/

Neuhaus, A. H., Urbanek, C., Opgen-Rhein, C., Hahn, E., Ta, T. M. T., Koehler, S., Gross, M., & Dettling, M. (2010). Event-related potentials associated with Attention Network Test. International Journal of Psychophysiology, 76(2), 72–79. 10.1016/j.ijpsycho.2010.02.005

Paavola, M. L., Mordkoff, J. T., & Moore, C. M. (2026). Simon says “stay in touch”: Reachability moderates the effect of irrelevant spatial congruence. Psychonomic Bulletin & Review, 33(4), Article 133. 10.3758/s13423-026-02879-7

Parsons, T. D. (2015). Virtual reality for enhanced ecological validity and experimental control in the clinical, affective and social neurosciences. Frontiers in Human Neuroscience, 9, Article 660. 10.3389/fnhum.2015.00660

Petersen, S. E., & Posner, M. I. (2012). The attention system of the human brain: 20 years after. Annual Review of Neuroscience, 35, 73–89. 10.1146/annurev-neuro-062111-150525

Pion-Tonachini, L., Kreutz-Delgado, K., & Makeig, S. (2019). ICLabel: An automated electroencephalographic independent component classifier, dataset, and website. NeuroImage, 198, 181–197. 10.1016/j.neuroimage.2019.05.026

Polich, J. (2007). Updating P300: An integrative theory of P3a and P3b. Clinical Neurophysiology, 118(10), 2128–2148. 10.1016/j.clinph.2007.04.019

Posner, M. I., & Petersen, S. E. (1990). The attention system of the human brain. Annual Review of Neuroscience, 13, 25–42. 10.1146/annurev.ne.13.030190.000325

Praamstra, P. (2007). Do’s and don’ts with lateralized event-related brain potentials. Journal of Experimental Psychology: Human Perception and Performance, 33(2), 497–502. 10.1037/0096-1523.33.2.497

Praamstra, P., & Oostenveld, R. (2003). Attention and movement-related motor cortex activation: A high-density EEG study of spatial stimulus-response compatibility. Cognitive Brain Research, 16(3), 309–322. 10.1016/S0926-6410(02)00286-0

Santhana Gopalan, P. R., Loberg, O., Hämäläinen, J. A., & Leppänen, P. H. T. (2019). Attentional processes in typically developing children as revealed using brain event-related potentials and their source localization in Attention Network Test. Scientific Reports, 9(1), Article 2940. 10.1038/s41598-018-36947-3

Simon, J. R., & Rudell, A. P. (1967). Auditory S-R compatibility: The effect of an irrelevant cue on information processing. Journal of Applied Psychology, 51(3), 300–304. 10.1037/h0020586

Tekampe, D., Sulaj, A., Hausfeld, L., & Schwartze, M. (2023). A virtual reality implementation of the Attention Network Test-Revised. In J. Balint and J. Fels (Eds.), Proceedings of the 1st AUDICTIVE Conference: June 19-22, 2023, RWTH Aachen University, Aachen, Germany (Vol. 1, pp. 142–145). RWTH Aachen University. 10.18154/RWTH-2023-08887

Tekampe, D. L., Sulaj, A., Tekampe, P., Hausfeld, L., & Schwartze, M. (2026). Attention networks and their interactions: Moving from the lab to mobile virtual reality. Heliyon, 12(1), Article e44267. 10.1016/j.heliyon.2025.e44267

Woodman, G. F., Arita, J. T., & Luck, S. J. (2009). A cuing study of the N2pc component: An index of attentional deployment to objects rather than spatial locations. Brain Research, 1297, 101–111. 10.1016/j.brainres.2009.08.011

