## Supplementary Materials for "Neural correlates of location–response compatibility in an immersive virtual-reality Attention Network Test: a multiverse electroencephalography analysis"

[1] University of Luxembourg, Department of Behavioural and Cognitive Sciences, Luxembourg
[2] Independent Researcher
[3] Maastricht University, Department of Neuropsychology & Psychopharmacology, the Netherlands
[4] Maastricht University, Department of Cognitive Neuroscience, the Netherlands

### Supplementary Materials

### S1. Supplementary Methods

#### S1.1. Preprocessing multiverse specification

The preprocessing multiverse was implemented as a restricted, balanced specification space rather than as an exhaustive search across all possible EEG pipelines. The 192 branches crossed high-pass filter, low-pass filter, line-noise attenuation, ASR policy, reference policy, ICLabel rejection policy, and cleaning-stringency bundle: 2 high-pass levels × 2 low-pass levels × 2 line-noise methods × 2 ASR policies × 2 reference policies × 3 ICLabel policies × 2 cleaning-stringency bundles = 192 branches.

High-pass filters were 0.1 or 0.5 Hz. Low-pass filters were 30 or 40 Hz. Line-noise attenuation was implemented as either CleanLine or a 50-Hz notch filter. ASR policy was either ASR enabled or ASR disabled. Reference policy was average reference or linked mastoids. ICLabel policy was conservative, moderate, or eye/muscle-only. Cleaning stringency was implemented as a composite bundle rather than as three independent decisions. The canonical bundle used a −200 to 0 ms target baseline, 100 µV epoch rejection, and no gyro-sensitive epoch rejection. The strict/gyro-sensitive bundle used a −100 to 0 ms target baseline, 75 µV epoch rejection, and movement-sensitive epoch rejection based on gyroscope magnitude.

The branch space was selected to represent preprocessing decisions that are common in ERP research and potentially consequential for lateralised EEG recorded during immersive VR. These included filtering, line-noise handling, reference selection, ASR, ICA-based artefact rejection, baseline window, amplitude rejection, and movement-sensitive epoch handling. Branches were interpreted as alternative defensible specifications applied to the same raw data, not as independent replications. The cleaning-stringency factor was interpreted as a composite processing decision; differences associated with this factor are therefore not attributed to baseline window, amplitude threshold, or gyro-sensitive epoch rejection in isolation.

Filtering was implemented in EEGLAB using zero-phase non-causal FIR filters. Branch-specific ERP-signal filters were applied at 0.1 or 0.5 Hz high-pass and 30 or 40 Hz low-pass settings. ICA-copy data were high-pass filtered at 1 Hz while otherwise following the branch-specific low-pass, line-noise, ASR, bad-channel exclusion, and reference settings. CleanLine branches used 50-Hz line-frequency attenuation implemented with the CleanLine EEGLAB plug-in (Mullen, 2012); notch branches used a 49-51 Hz band-stop filter. Exact software calls, filter orders, transition bandwidths, and −6-dB cut-offs are documented in the analysis code and branch audit exports.

#### S1.2. Bad-channel handling, ASR, ICA, and ICLabel

Raw EEG was imported from XDF files. Channel locations were assigned to recorded EEG channels, and FCz was appended as a channel-location/reference label rather than treated as a recorded electrode. Gyroscope channels were retained for movement auditing but excluded from EEG-channel preprocessing, feature extraction, and statistical analyses.

Bad-channel handling followed a staged conservative policy. The prespecified T7 decision and automatically detected bad channels were logged for each branch × participant dataset. Automatic bad-channel detection used channel-wise standard deviation, robust z scores based on the median and median absolute deviation, and maximum absolute amplitude across EEG channels. Bad channels were temporarily neutralised for ASR, when necessary, excluded from pre-ICA referencing, ICA estimation, and epoch artefact rejection, and interpolated only after IC removal. Datasets failed preprocessing QC if more than 20% of EEG channels would require interpolation or if C3 or C4 was marked for interpolation.

When selected by the branch, ASR was applied using the implementation provided in the EEGLAB *clean_rawdata* plug-in. Channels marked as bad were temporarily interpolated for the ASR operation; after ASR, only channels not marked as bad were copied back into the working EEG dataset. ASR audit rows recorded the ASR policy, requested burst criterion, pre-ASR RMS amplitude, post-ASR RMS amplitude, RMS change, and any fallback note. When ASR was disabled, no ASR correction was applied.

ICA was estimated on a 1-Hz high-pass copy of the branch-specific continuous data. ICA channel indices excluded bad channels and appended FCz. The PCA dimension was determined from the effective-rank audit and bounded by the number of usable ICA channels and a maximum of 22. Extended runica was used. ICA weights were transferred to the ERP-signal data, components were classified using ICLabel, and components were removed according to the branch-specific ICLabel policy. The conservative policy rejected eye, muscle, heart, channel-noise, and line-noise components at probability ≥ .90. The moderate policy rejected the same classes at probability ≥ .80. The eye/muscle-only policy rejected only eye and muscle components at probability ≥ .80. ICLabel classifications for linked-mastoid branches were interpreted cautiously because

ICLabel was developed primarily for components standardised to common-average reference.

#### S1.3. Event decoding, epoching, quality checks (QC), SME, and movement diagnostics

ANT-VR event markers encoded trial phase, cue condition, cue-target interval, target location, target identity, target-defined response side, flanker congruency, and button responses. The pipeline decoded markers into branch-specific trial tables and epoch metadata. Event-timing checks evaluated trial-phase order, cue-target interval coding, target-to-response timing, duplicated or missing response markers, and consistency between decoded trial tables and epoch metadata. Target events were used as the temporal anchor for target-locked epochs and observed button responses were used as the temporal anchor for response-locked epochs.

Target-locked epochs spanned −500 to +1000 ms around target onset. Response-locked epochs spanned −800 to +500 ms around the observed response. Target-locked baseline correction used either −200 to 0 ms or −100 to 0 ms according to the branch-specific cleaning-stringency bundle. Response-locked baseline correction used −400 to −200 ms in all branches. Epoch-level metadata were mandatory for every analysed EEG file; any mismatch between EEG epoch count and metadata rows was treated as a fatal analysis error.

Standardised measurement error was computed from branch-specific epoch amplitudes as the standard deviation of epoch-level mean amplitudes divided by the square root of the number of epochs. SME was exported for target-locked P1, posterior N1, N2pc ROI quality, frontocentral N2, centroparietal P3, parietal P3, C3/C4 lateralisation quality windows, and response-locked C3/C4 windows. SME rows were summarised overall and by location-response compatibility, response side, flanker congruency, and cue-target interval.

Movement and C3/C4 artefact diagnostics were exported for target-locked and response-locked streams. Target-locked diagnostic windows included 0-600, 300-600, 250-500, and 350-650 ms. Response-locked diagnostic windows included −300 to +100, −200 to 0, −100 to +100, and 0 to +200 ms. For each window, the pipeline recorded mean absolute C3 amplitude, mean absolute C4 amplitude, mean absolute C3-minus-C4 amplitude, mean absolute first differences for C3 and C4, gyro-flag rate, and high-frequency proxy availability. These diagnostics were interpreted as quality-context summaries rather than as optimisation criteria for selecting a preferred preprocessing branch.

#### S1.4. Secondary, sensitivity, and falsification analyses

Secondary moderator analyses tested whether the compatibility contrast varied with flanker congruency, spatial cue validity, or cue-target interval. Strict moderator models retained the balanced structure used for the compatibility contrast and were interpreted as secondary to the main compatibility contrast. Collapsed moderator models used less sparse cell structures and were not treated as substitutes for the balanced model.

Response-locked C3/C4 analyses used the independently generated response-locked stream and estimated incompatible-minus-compatible effects in the −200 to 0 ms, −100 to +100 ms, and 0 to +200 ms windows around response onset. C3/C4 decomposition analyses recomputed target-locked C3/C4 activity between 300 and 600 ms relative to response side, target-defined response side, target location, and raw C4-minus-C3 amplitude. Trial-threshold sensitivity analyses repeated target-family estimates using minimum cell thresholds of 1, 3, 6, 8, and 10 trials.

Pre-target negative-control analyses used the −200 to 0 ms target-locked window. Sign-exchange diagnostics evaluated whether the observed median branch effect and directional stability differed from a participant-level sign-exchange null. Participant-resampling bootstrap uncertainty was estimated with 1,000 bootstrap resamples across the 192 preprocessing branches. Repeated stratified split-half reliability was estimated with 100 repetitions and Spearman-Brown correction for the key lateralised target families. These controls were interpreted as falsification, uncertainty, reliability, or sensitivity analyses rather than as independent replications of the target-locked N2pc compatibility effect. Holm correction was applied separately within the secondary diagnostic families: the N2pc and response-referenced C3/C4 pre-target controls, the four target-locked C3/C4 reference-frame measures, the three C3/C4 pre-target decomposition controls, and the three response-locked C3/C4 windows.

### S2. Supplementary Results

#### S2.1. Multiverse flow and preprocessing quality checks

The multiverse contained 192 preprocessing branches and 44 participants in the final behavioural-overlap sample. Across branches, 9,024 branch × participant preprocessing requests were launched. A total of 8,676 requests generated a quality-control row, including 7,849 QC-pass rows and 827 QC-fail rows. The remaining 348 requests terminated before QC-row generation because linked-mastoid reference was requested when M1 or M2 had been marked bad in the corresponding preprocessing request. These rows were treated as pre-QC attrition rather than preprocessing QC failures.

After QC-pass filtering and restriction to the behavioural-overlap sample, 7,369 target-locked and 7,369 response-locked branch × participant datasets entered feature extraction. Analysed participants per branch ranged from 30 to 42. The median number of correct retained target-locked epochs per branch was 10,101, IQR [9,438, 10,846], and the median target-locked retention rate was 95.5%, IQR [95.3%, 95.7%]. The channel-label audit found exactly one C3 and one C4 label in every analysed target-locked and response-locked dataset.

#### S2.2. Full branch-level multiverse outputs

Full branch-level outputs report all 192 preprocessing branches, branch-level participant counts, incompatible-minus-compatible estimates, standard deviations, confidence intervals, *t* statistics, degrees of freedom, *p* values, Holm-adjusted *p* values, FDR-adjusted *p* values, Cohen’s dz, and preprocessing-decision labels for each target family. The branch-level tables include P1, posterior N1, N2pc, frontocentral N2, centroparietal P3, parietal P3, response-referenced C3/C4, target-location-referenced C3/C4, target-defined response-side C3/C4, raw C4-minus-C3, and response-locked C3/C4 windows where applicable. These rows are interpreted as dependent robustness summaries rather than independent replications.

#### S2.3. Behavioural-stream and EEG-retention checks

Full-stream behavioural checks used the complete 1,700-ms response window. Location-response incompatibility was not associated with a reliable reaction-time difference, mean difference = +2.65 ms, 95% CI [−5.97, +11.27], $t(43) = 0.62$, $p = .539$, or accuracy difference, mean difference = −0.76 percentage points, 95% CI [−2.08, +0.56], $t(43) = -1.16$, $p = .254$. Late-response rates were low in both compatibility conditions and did not differ reliably at participant level. Across branches, incompatible trials showed a small median reduction in EEG retention relative to compatible trials. These behavioural and retention analyses were treated as EEG-selection diagnostics rather than as a re-analysis of the complete behavioural ANT-VR study.

#### S2.4. Diagnostic controls and sensitivity analyses

Pre-target negative-control analyses in the −200 to 0 ms target-locked window did not reproduce the target-locked N2pc or target-location C3/C4 pattern. Response-locked C3/C4 analyses showed positive median effects in windows before, around, and after response onset but no Holm-corrected branch-level support. Sign-exchange diagnostics supported the observed N2pc and target-location C3/C4 median effects relative to the participant-level sign-exchange null ($p$ = .034 and $p$ = .010, respectively), but not the corresponding directional-stability statistics ($p$ = .180 and $p$ = .265, respectively), whereas raw C4-minus-C3 did not show comparable evidence.

Trial-threshold sensitivity analyses repeated target-family estimates using minimum cell thresholds of 1, 3, 6, 8, and 10 trials. The median N2pc-window posterior-lateralisation contrast remained positive across thresholds, although the proportion of branches with an unadjusted $p < .05$ varied. Target-location C3/C4 remained positive in all branches at every threshold. Raw C4-minus-C3 remained close to zero. Response-referenced C3/C4 varied more strongly across trial-count thresholds. These analyses indicate that the direction of the median N2pc-window posterior-lateralisation contrast did not depend on the selected minimum cell threshold, whereas response-referenced C3/C4 was more threshold-sensitive.

### S2.5. Measurement quality, preprocessing-decision sensitivity, bootstrap uncertainty, and reliability

Standardised measurement error, movement-proxy, ASR RMS-change, bootstrap, and split-half reliability analyses were used to contextualise the multiverse results. The N2pc ROI-quality window showed a median SME of 0.278 µV, IQR [0.258, 0.309]. Target-locked C3/C4 and response-locked peri-response C3/C4 showed comparable SME ranges. Incompatible trials showed only a very small SME increase relative to compatible trials. Across branch × participant observations, N2pc magnitude was only weakly associated with mean SME and was not materially associated with the C3/C4 movement proxy. ASR audit rows showed a median ASR-associated RMS change of −7.02 µV, IQR [−11.17, −4.20].

Preprocessing-decision sensitivity summaries indicated that the largest descriptive median differences in the N2pc estimate were associated with ASR policy, the cleaning-stringency bundle, and the high-pass filter. Smaller descriptive differences were observed for low-pass filter, line-noise attenuation, reference policy, and ICLabel policy. These summaries were not analysed as independent observations and should not be interpreted as causal effects of preprocessing decisions or as evidence for a single optimal branch.

Participant-resampling bootstrap uncertainty supported a positive median branch effect for N2pc and target-location C3/C4, whereas response-referenced C3/C4 and raw C4-minus-C3 bootstrap intervals included zero. Split-half reliability was weak for N2pc, indicating that the N2pc-window posterior-lateralisation contrast should be interpreted as a group-level experimental effect rather than as an individual-difference measure or biomarker candidate. Target-location and response-referenced C3/C4 showed modest split-half reliability but remained secondary reference-frame analyses rather than replacements for the N2pc endpoint.

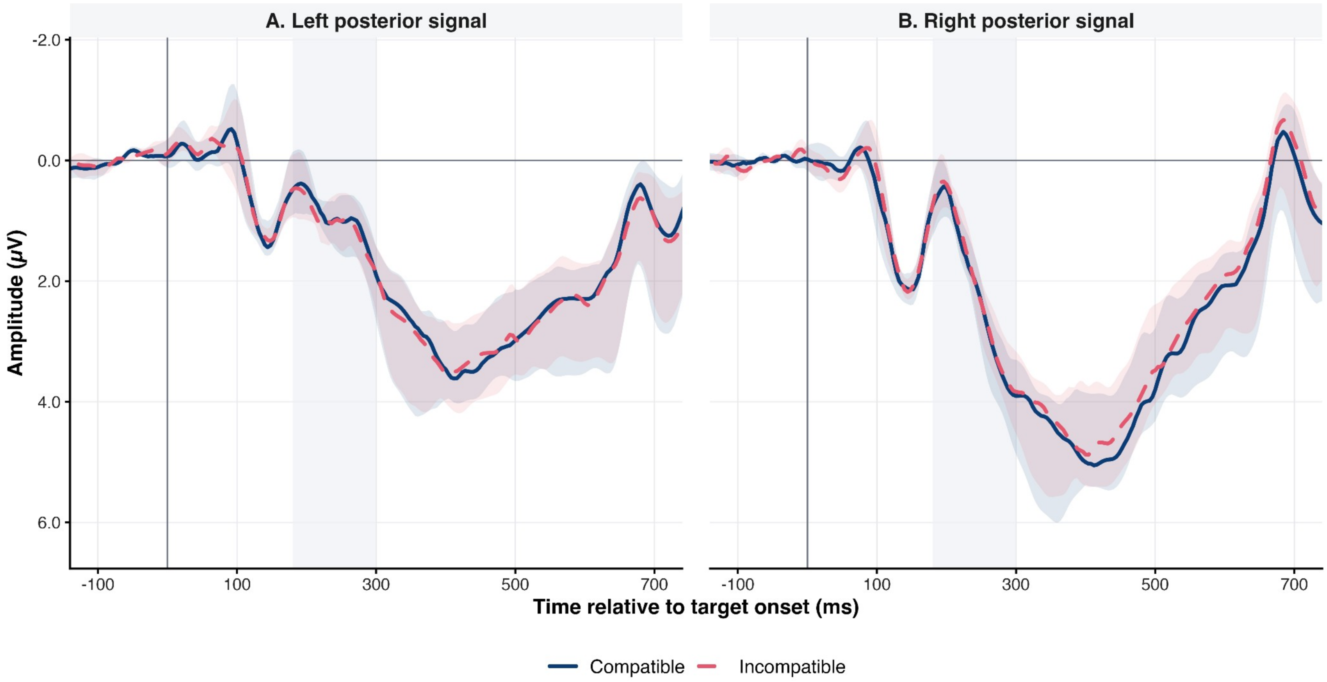


**Supplementary Figure S1.** Constituent left- and right-posterior waveforms by location-response compatibility

**Note.** Waveforms show the posterior signals underlying the compatibility contrasts presented in Figure 2. Panel A shows the left-posterior P7 signal separately for compatible and incompatible trials, and Panel B shows the corresponding right-posterior P8 signal. Within each preprocessing branch, participant waveforms were averaged using retained trial counts as weights; the displayed lines represent the median across the 192 preprocessing specifications. Shaded ribbons indicate the interquartile range across preprocessing branches. The grey band marks the prespecified N2pc analysis window from 180 to 300 ms. These waveforms are presented descriptively; the inferential analysis was based on the incompatible-minus-compatible contrasts shown in Figure 2.

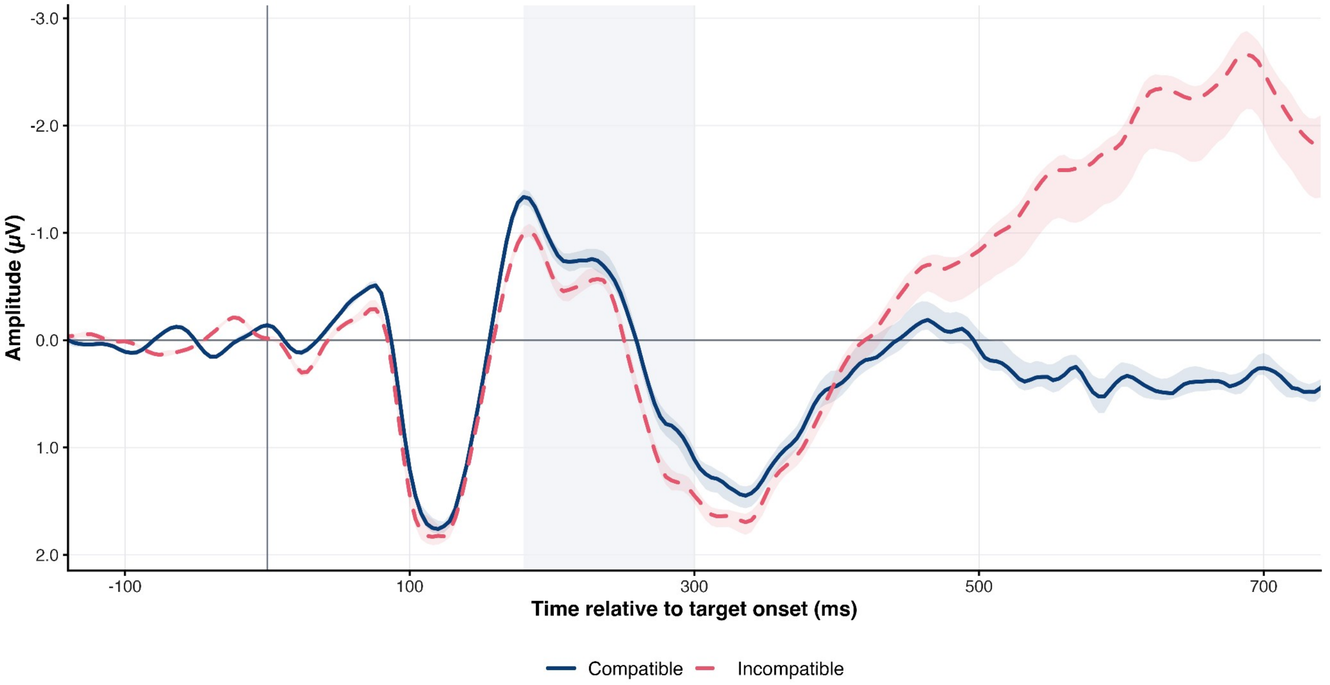


**Supplementary Figure S2.** N2pc Waveform Diagnostic

**Note.** Lines show median branch-level N2pc contra-minus-ipsilateral waveforms for compatible and incompatible trials. Shaded ribbons indicate interquartile ranges across preprocessing branches, and the grey band marks the N2pc analysis window. Waveforms are descriptive diagnostics; inferential analysis is based on prespecified mean-amplitude contrasts.

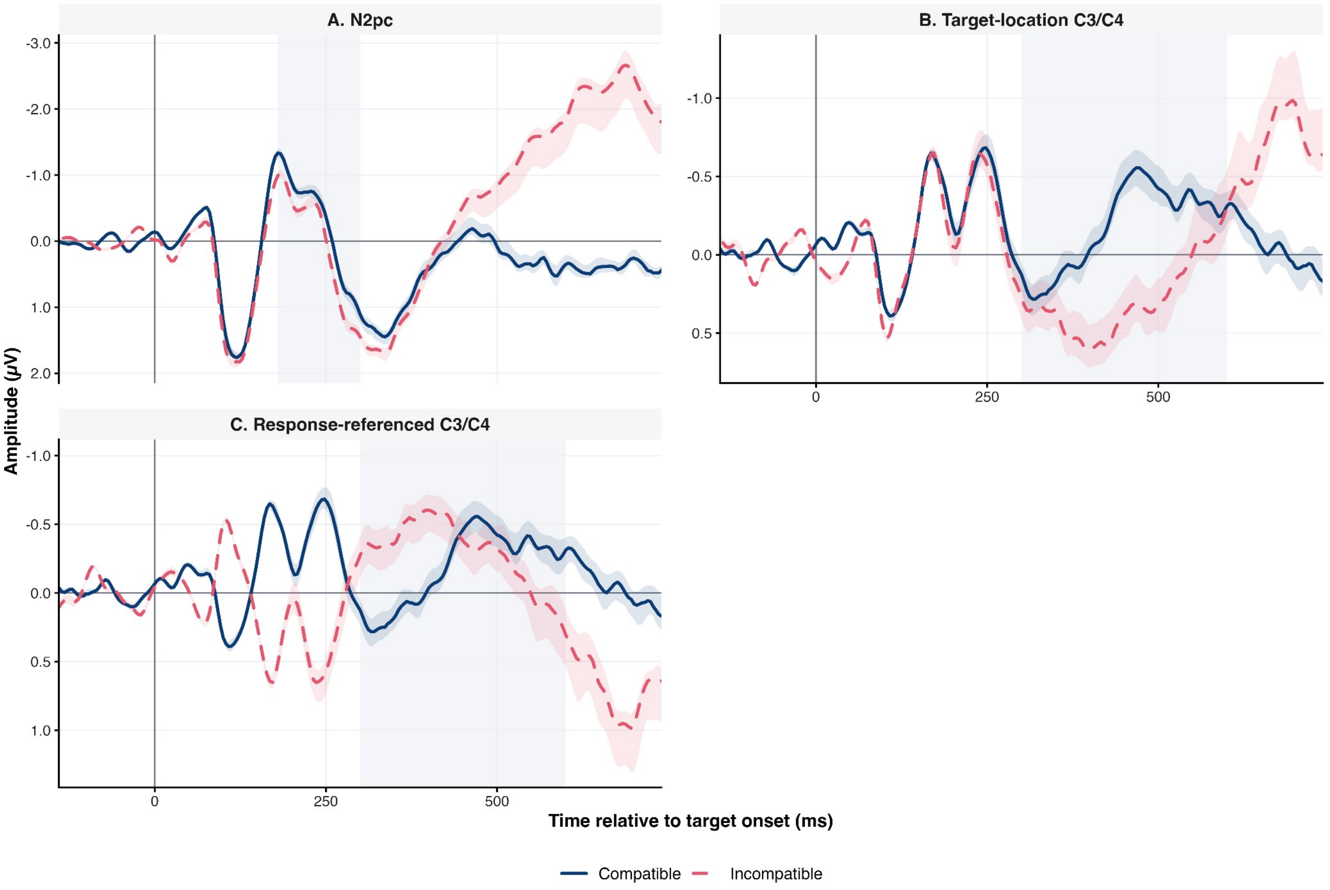


**Supplementary Figure S3.** Reference-frame comparison waveforms

**Note.** Lines show median branch-level waveforms for N2pc, target-location-referenced C3/C4, and response-referenced C3/C4. Shaded ribbons indicate interquartile ranges across preprocessing branches, and grey bands mark the corresponding analysis windows. These waveform panels provide descriptive context for the branch-level secondary reference-frame analyses.
